# Discovery and structure of a widespread bacterial ABC transporter specific for ergothioneine

**DOI:** 10.1101/2022.05.02.490363

**Authors:** Yifan Zhang, Giovanni Gonzalez-Gutierrez, Katherine A. Legg, Katherine A. Edmonds, David P. Giedroc

**Author notes:** These authors share senior authorship (Katherine A. Edmonds, David P. Giedroc).

## Abstract

Ergothioneine (ET) is the 2-thiourea derivative of trimethylhistidine that is biosynthesized only by select fungi and bacteria, notably *Mycobacterium tuberculosis*, and functions as a potent scavenger of reactive oxygen species. Although ET is obtained in the diet and accumulates in vertebrate cells via an ET-specific transporter, the extent to which ET broadly functions in bacterial cells unable to synthesize it is unknown. Here we show that *spd_1642-1643* in *Streptococcus pneumoniae* D39, a Gram-positive respiratory pathogen, encodes a novel ergothioneine uptake ATP-binding cassette (ABC) transporter, which we designate EgtUV. EgtU is a permease-solute binding domain (SBD) fusion protein, and the SBD binds ET with high affinity and exquisite specificity in the cleft between the two subdomains, with cation-π interactions engaging the betaine moiety and a water-mediated hydrogen bonding network surrounding the C2-sulfur-containing imidazole ring. Bioinformatics studies reveal that EgtUV is uniquely strongly conserved among known quaternary amine-specific transporters and widely distributed in firmicutes, including the human pathogens *Listeria monocytogenes*, as BilEB, *Enterococcus faecalis* and *Staphylococcus aureus*. This discovery significantly diversifies the LMW thiol pool in Gram-positive human pathogens that may contribute to antioxidant defenses in the infected host.

## MAIN

Cell-abundant low molecular weight (LMW) thiols maintain the reducing environment of the cytoplasm of bacterial cells and the cytosol of eukaryotic cells, and include the ubiquitous tripeptide glutathione^1^. Bacteria unable to access glutathione synthesize other thiols, including bacillithiol and mycothiol found in some firmicutes and actinomycetes, respectively^2,3^ or other, inorganic molecules such as thiosulfate^4^. These cell-abundant LMW thiols provide protection against endogenous or exogenous reactive oxygen and nitrogen species (ROS, RNS) as ROS scavengers to create thiol disulfides, which are reduced by NADPH-dependent oxidoreductases, regenerating the free thiol and thus maintaining redox balance^5^. Bacterial LMW thiols are known to play important roles in oxidative and reductive stress responses in the infected host^6-8^.

Ergothioneine (ET) is a LMW thiol and trimethylamine (betaine) derivative of histidine with a thione group installed at the imidazole C2 position^9^. Unlike other LMW thiols that function as cellular redox buffers, ET is primarily found in the thione tautomer rather than the thiol tautomer at physiological pH^10,11^. This thiol-thione tautomerization significantly increases (makes more positive) the thiol-disulfide reduction potential of ET relative to other LMW thiols (*E*º of –0.06 vs – 0.28 V) ^12-17^. This endows ET with properties of a highly effective scavenger of myriad ROS, including host-derived oxidants HOCl, peroxynitrite (ONOO^−^), and nitrosoperoxycarbonate (ONOOCO_2_^−^), due to its intrinsic resistance to autoxidation^18-20^. ET, like other LMW thiols^21^, is able to chelate divalent metals including Cu^2+^ and Fe^2+^ forming 2:1 ligand:metal complexes^22,23^.

ET biosynthesis has been reported to occur only in select filamentous fungi including *Neurospora crassa*^24^, certain cyanobacteria^25^, Methylobacterium *spp*.^26^, Burkholderia *spp*.^27^, and in actinomycetes, including the causative agent of tuberculosis, *Mycobacterium tuberculosis*, and its soil saprophyte, *Mycobacterium smegmatis*^28-30^. ET biosynthesis generally begins with *L*-histidine, which is trimethylated by a SAM-dependent α-amine methyltransferase into *L*-hercynine, with the sulfur atom derived from a LMW thiol donor, via the a hercynyl-thiol sulfoxide intermediate.^31-34^ Recent work reveals that ET can also be catabolized by soli-dwelling organisms, as well as by GI tract organisms, likely impacting ET bioavailability in those niches^35^.

The physiological functions of endogenous ET in various bacterial and fungal organisms have been extensively investigated^36^. In mycobacteria, ET and mycothiol are the two major LMW thiol buffers, and although ET has a much lower cellular abundance relative to mycothiol, it specifically functions in ROS detoxification^37,38^. In *M. tuberculosis*, ET maintains bioenergetics homeostasis when fatty acids are a major carbon source, while also modulating the susceptibility toward anti-tuberculosis drugs^37,39,40^. ET is also important for the survival of *M. tuberculosis* in macrophages and for virulence in mouse models of infection^41^. In *N. crassa*, ET is a major antioxidant that protects the organism against peroxide damage during germination^42,43^. In *Aspergillus fumigatus*, ET functions as an antioxidant that provides protection from free radicals generated by oxidative stress, while also functioning in the response to transition metal toxicity, including iron, zinc, copper and cobalt^44,45^. In the fission yeast, *Schizosaccharomyces pombe*, peroxide stress and Cd^2+^ stress also stimulate ET biosynthesis^46^.

There have been no reports of ET biosynthesis in plants or animals^36,47^. In humans, ET is obtained from the diet and accumulates in tissues by an ergothioneine specific transporter ETT, previously named organic cation/carnitine transporter I (OCTN1)^48-50^. The expression level of ETT in various tissues has been used as proxy for abundance and distribution of ET in animals^47,48,51^. ETT expression is high in the small intestine and the kidney which reflects dietary ET uptake and ET recovery from the urine, respectively^49,52,53^. High ETT expression has also been identified in blood cells, including erythrocytes in bone marrow, granulocytes, monocytes and neutrophils, while detectable expression occurs in other tissues including the lung^20,49,52,54,55^ Decreased blood or plasma levels of ET have been used as a biomarker in many human diseases and disorders, consistent with the description of ET as a nutritional “antioxidant” that provides defense against ROS or RNS that are generated during injury or disease^47^.

These studies suggest that ET is significantly bioavailable in vertebrates and could be exploited by both resident commensals and pathogenic organisms to provide protection against host oxidative insults; however, no widespread bacterial transporter for ET is known^36,47,48^. Here, we describe the discovery and structural characterization of a bacterial ET transporter. We show that genes *spd_1642-1643* in the Gram-positive commensal and respiratory pathogen, *Streptococcus pneumoniae* D39, encode a novel prokaryotic ET uptake ABC transporter, which we denote EgtUV. Quantitative LMW thiol profiling reveals that EgtUV is required for ET accumulation in *S. pneumoniae*. The ET-bound crystal structure of the solute binding domain (SBD) of EgtU, coupled with extensive NMR studies, provide novel insights into ET affinity, binding mechanism and specificity. Bioinformatics analyses and accompanying biophysical studies reveal that EgtUV is widely distributed in firmicutes including the human pathogens *Enterococcus faecalis, Staphylococcus aureus* and *Listeria monocytogenes*, the latter as a bile acid exclusion and virulence determinant^56^. This discovery dramatically expands the diversity of the LMW thiols to include ET in an important human pathogen, well beyond *M. tuberculosis*, where it may contribute to antioxidant defenses in the infected host.

## RESULTS

### *Spd_1642-1643* encodes an ABC transporter specific for ET

We recently identified an uncharacterized operon in *Streptococcus pneumoniae* D39, *spd_1642-1645*, that is highly conserved in *Streptococci* and is regulated in part by a quinone-sensing Rrf2 family transcriptional regulator, SifR^57^. The SifR regulon allows access to a host-derived nutritional catechol-iron source, while avoiding oxidative stress associated catechol oxidation^57^. This operon encodes an uncharacterized MarR family transcriptional regulator^58^ (*spd_1645*), a putative Snoal2 family polyketide cyclase/hydrolase^59^ (*spd_1644*) and an ABC transporter annotated as an osmoprotectant uptake (Opu or Pro) system that transports quaternary amines, *e*.*g*., glycine betaine^60^ (*spd_1642-1643*) (Fig. 1a). Given the SifR connection to redox stress and iron assimilation, we hypothesized that SPD_1642 is involved in ET uptake and thus provisionally named this gene *egtU* (<u>e</u>r<u>g</u>othioneine <u>u</u>ptake).

**Figure 1.**
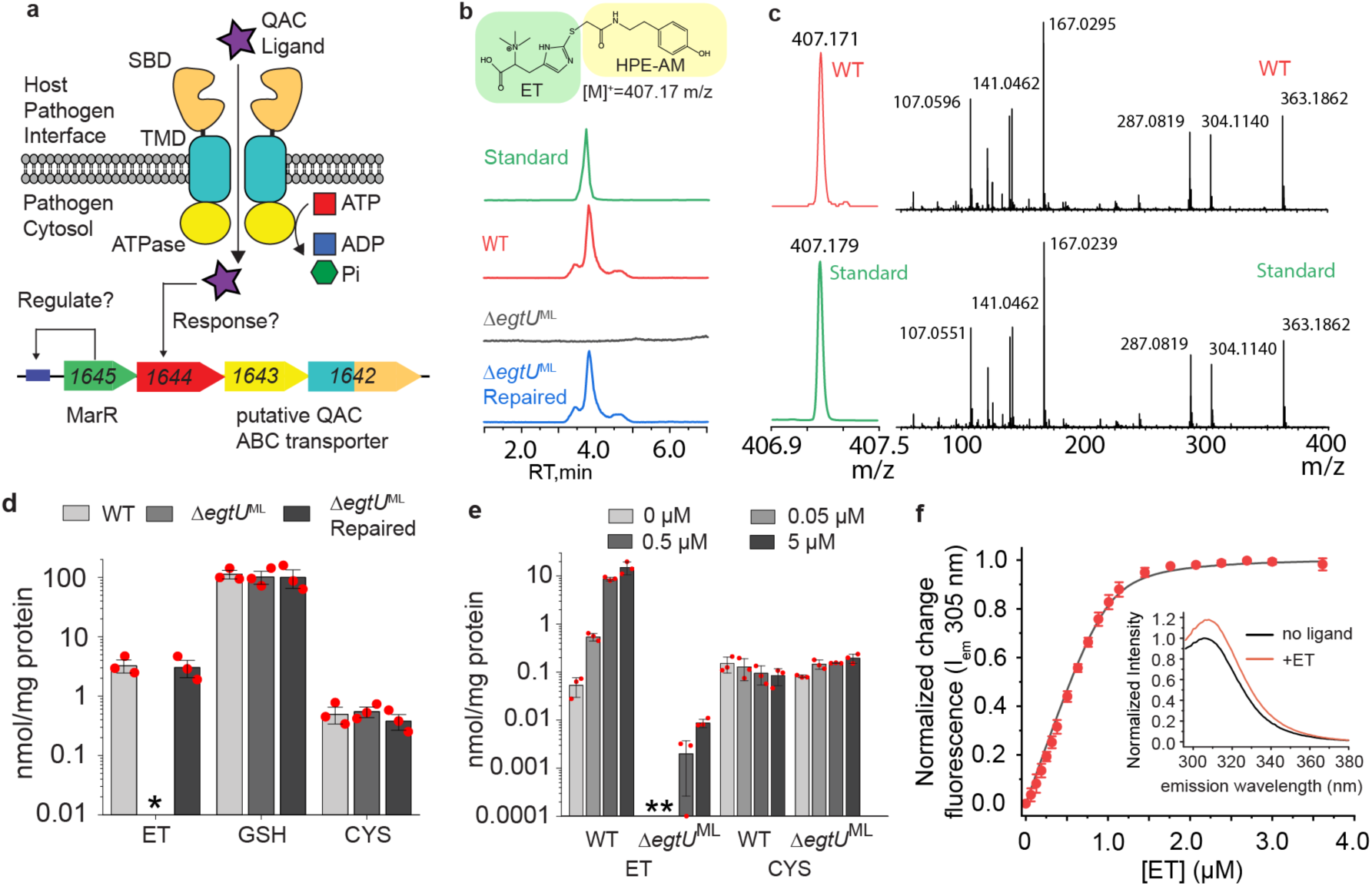
*spd_1643-1642* encodes an ET uptake transporter in *S. pneumoniae* denoted EgtUV. **a**, Hypothesized functional roles of the genes of the streptococcal conserved operon^57^ harboring a candidate QAC ABC transporter, encoded by *spd_1642-1643*. **b**, Structure of HPE-IAM-derivatized (capped) ET and LC traces of capped ET from authentic ET (standard) and cell lysates from the indicated *S. pneumoniae* D39 strains grown on BHI media. WT, wild-type; Δ*egtU*^ML^, markerless (ML) deletion of most of the *egtU* gene, which was then repaired via insertion of a wild-type *egtU* allele (Δ*egtU*^ML^ repaired). **c**, MS1 (*left*) and LC-MS/MS (*right*) spectra of HPE-IAM capped ET found in a wild-type (WT) cell lysate from cells grown in BHI vs. authentic ET (standard). **d**, Cellular concentrations of ET, GSH and CYS found in the indicated strains of *S. pneumoniae* D39 grown in BHI in biological triplicate with individual measurements shown by red filled circles. *, not detected (≤0.0001 nmol/mg protein). **e**, Cellular concentrations of ET and CYS in the indicated strains of *S. pneumoniae* D39 grown in CDM supplemented with indicated concentration of ET in biological triplicate. *, not detected. Individual measurements are shown as red filled circles. GSH is not detected in these cell lysates (≤0.0001 nmol/mg protein). The small amount of ET detected in the Δ*egtU*^ML^ strain is likely derived from residual ET that remains after extensive washing of cell pellets. **f**, titration of ET into *Sp*EgtU SBD monitored by a change in the Tyr fluorescence emission intensity. *Inset*, tyrosine emission spectra of *Sp*EgtU SBD in the absence (black) and presence (red) of saturating ET.

To test this idea, we used an isotope dilution mass spectrometry-based thiol profiling strategy to quantify LMW thiols present in lysates derived from exponentially growing *S. pneumoniae* cells (Fig. 1b,c; Supplementary Figures 1-4). We find that glutathione (GSH)^61^, cysteine (Cys) and ET are major LMW thiols in a wild-type *S. pneumoniae* D39 strain cultured in a brain-heart infusion (BHI) growth medium (Fig. 1d). Moreover, a markerless Δ*egtU* strain lacks detectable ET, while ET levels are restored in an *egtU*-repaired strain (Fig. 1d). In strong contrast, cellular levels of GSH and cysteine are unaffected by the loss of *egtU*. We next quantified pneumococcal thiol levels when grown in a chemically defined medium to which variable ET was added (0.05 to 5 µM). These studies reveal a concentration-dependent increase in cellular ET that is lost in the Δ*egtU* strain, with no impact on Cys levels and no detectable GSH (Fig. 1e). We therefore rename the transmembrane permease domain-solute binding domain (TMD-SBD) fusion protein SPD_1642 as EgtU and the ATPase SPD_1643 as EgtV, with EgtUV an ergothioneine-specific uptake transporter in *S. pneumoniae*.

### The EgtU SBD binds ET with high affinity

The intrinsic tyrosine fluorescence of purified EgtU SBD (residues 233 to 506) increases upon addition of ET, accompanied by a slight red shift in the emission spectrum (Fig. 1f). These data confirm that EgtU SBD binds ET as a 1:1 complex, and reveal a *K*_a_ of ≈2.0×10^7^ M^-1^ (Table 1). The perturbation of the tyrosine fluorescence suggests that EgtU may engage ET by trimethylamine cation-π interactions in the same manner as SBPs specific for osmoprotectants like glycine betaine and choline^60,62^.

**Table 1.**
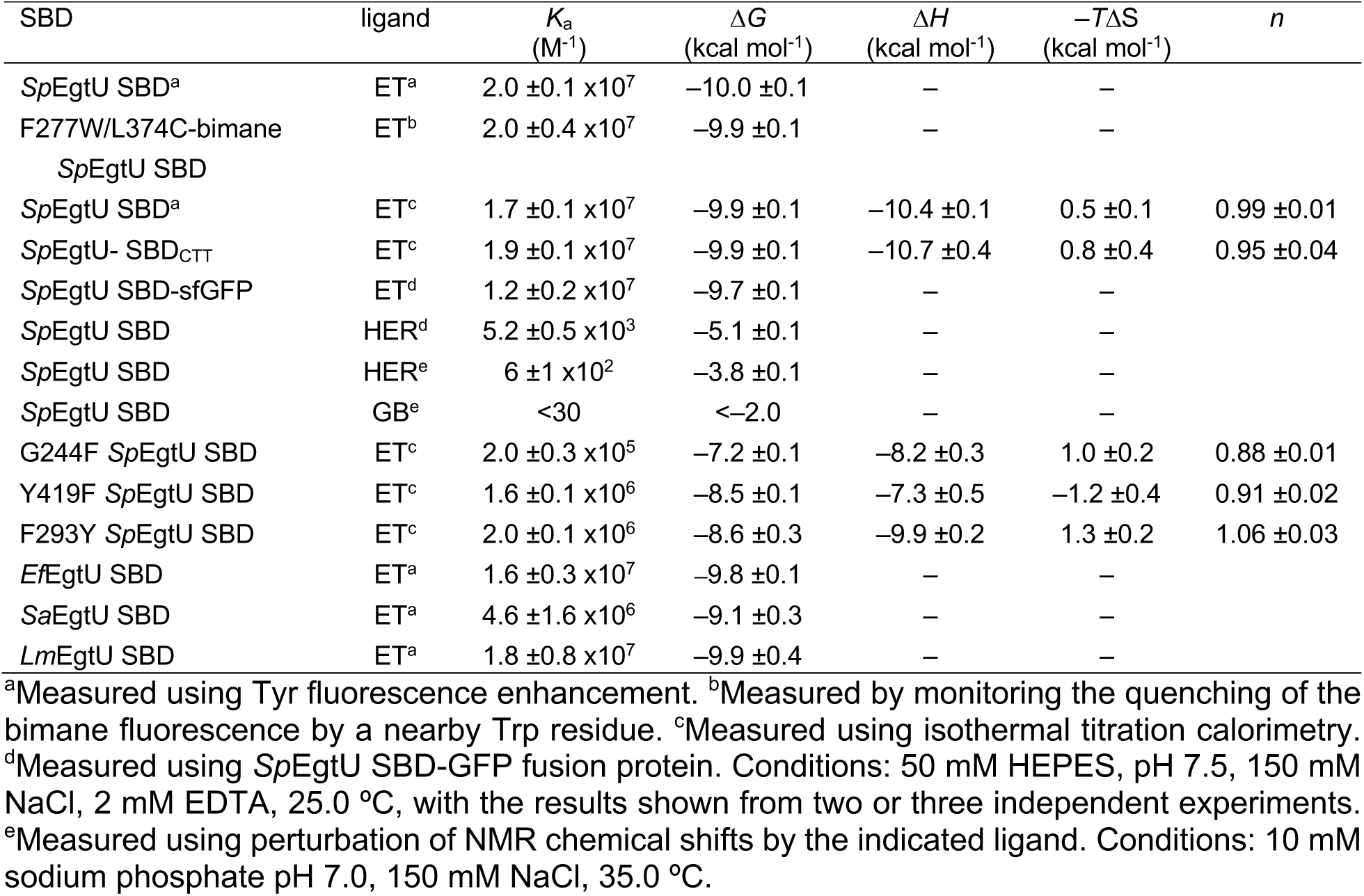
Summary of parameters obtained for the binding of ET and other ligands to wild-type and mutant *Sp*EgtU SBD.

### Crystal structure of the EgtU SBD-ET complex

To identify the molecular determinants of ET specificity in the *Sp*EgtU SBD, we determined the atomic structure using X-ray crystallography. Two structures of *Sp*EgtU SBD-ET complexes were independently obtained, at 1.82 and 2.44 Å resolution (Supplementary Table 1). The 1.82 Å structure of the holo-EgtU SBD, termed EgtU SBD_CTT_, harbors a short C-terminal truncation while the 2.44 Å structure is the wild-type domain. Isothermal titration calorimetry shows that EgtU SBD_CTT_ has ET binding affinity and thermodynamics that are identical to the wild-type protein (Supplementary Figure 5a, Table 1). The structures are virtually identical with a pairwise heavy-atom RMSD of 0.206 Å over the common regions (residues 233-501; Supplementary Figure 5b), and we therefore use the higher resolution EgtU SBD_CTT_-ET structure to describe its features. The structure of the EgtU SBD consists of two globular subdomains connected by a hinge consisting of two unstructured strands ≈10 residues long, with the ligand bound in the cleft between subdomains. The “terminal domain” encompasses the N-terminal residues 233-331 and C-terminal residues 445-506, while the “middle domain” consists of residues 341 to 432 (Fig. 2a). Each domain is characterized by a five-stranded β-sheet surrounded by five or six α-helices. The additional electron density found in the cleft between two domains corresponds precisely to that of *L*-ET (Fig. 2b), with the quaternary amine oriented toward the hinge and the bulky sulfur atom close to opening of the binding pocket.

**Figure 2.**
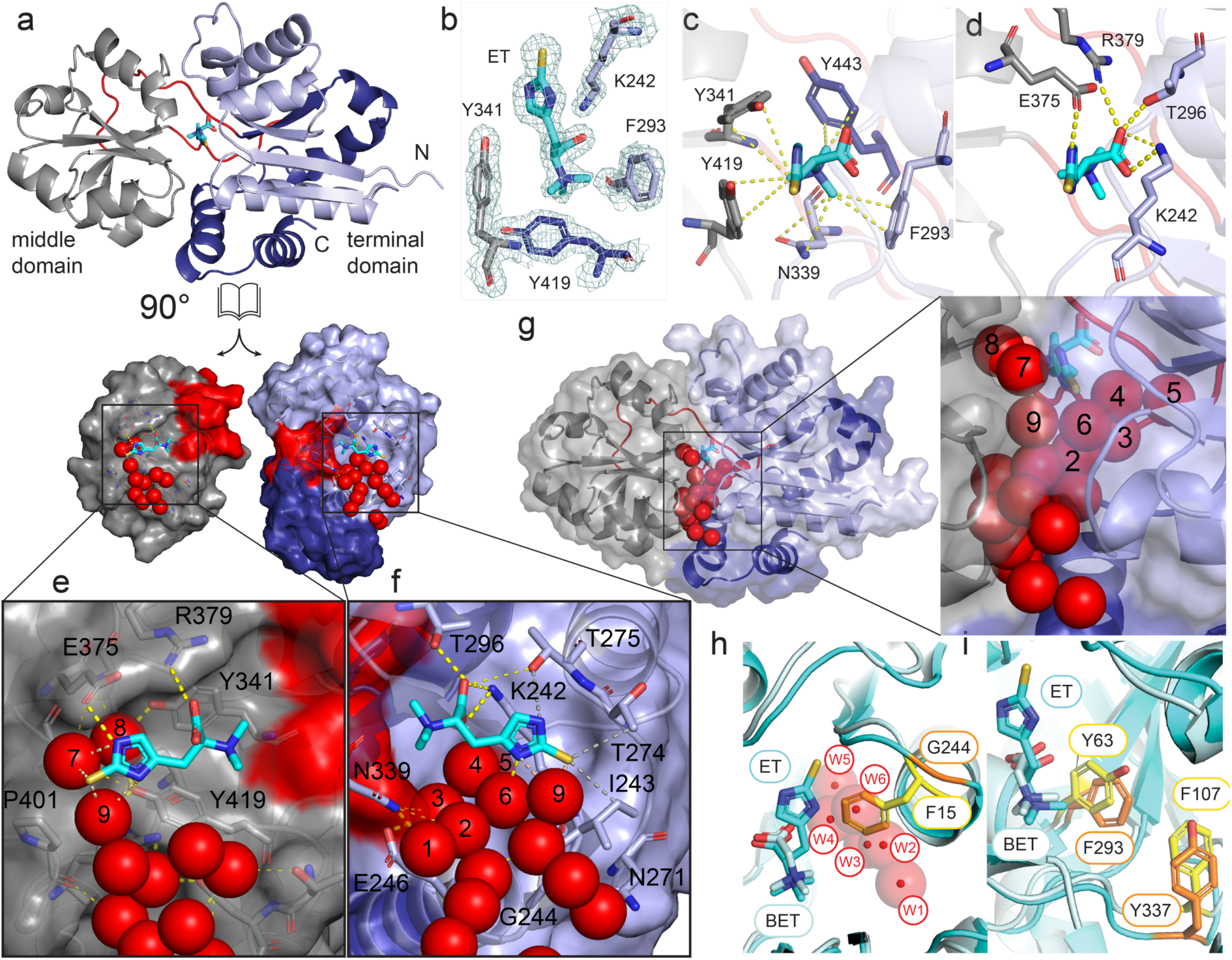
Structure of ET-bound *Sp*EgtU SBD. **a**, Crystal structure of ET-bound EgtU SBD_CTT_ shown as ribbon, with the terminal domain shaded light blue (residues 232-331) and deep blue (445-503), middle domain shaded gray (340-432) and domain linkers colored red. ET is shown as cyan sticks. **b**, Electron density map of ET and surrounding residues in ligand binding pocket. **c**, Quaternary amine region of the ET binding pocket, with residues in the aromatic pentagon shown as sticks, with polar and cation-π interactions shown as yellow dashed lines. **d**, Middle of the ET binding pocket, with polar interactions shown as gold dashed lines and interacting side chains shown as sticks. **e**, Thione sulfur region of the middle domain ET binding pocket, with surface colored as in panel **a**, side chains shown as sticks, water molecules as red spheres, and polar and hydrophobic interactions shown as yellow dashed lines. **f**, Thione sulfur region of the terminal domain ET binding pocket displayed as in panel **e. g**, High-occupancy water molecules (a subset are labeled 1-9 in panels **e, f** and **h**; see Supplementary Table 2) lining the binding pocket and interdomain cleft, shown as red spheres. **h**, Overlay of ligand binding pockets from the betaine SBP *Af*ProX (cyan) and ET SBP *Sp*EgtU SBD (palecyan), showing that a conserved G244 in *Sp*EgtU SBD provides space for a chain of water molecules (red spheres). **i**, Another view of these two ligand binding pockets that highlights the position of the F-Y switch between EgtU and an SBP specific for glycine betaine, with F293 and Y337 in *Sp*EgtU SBD and Y63 and F107 *Af*ProX shown as sticks.

The long, two-stranded hinge between domains identifies the EgtU SBD as a type II SBP^63,64^, and more specifically as a member of cluster F^65^. The EgtU SBD belongs to subcluster F-III, which employs a conserved set of aromatic residues arranged in a cage-like structure to coordinate the quaternary amine of the histidine betaine moiety. As in other subcluster F-III quaternary ammonium compound (QAC) binding proteins, notably *Archaeoglobus fulgidus* ProX^66^, the aromatic residues Y341, Y419, Y443 and F293 form four sides of a pentagon and contribute cation-π interactions, while the base of the pentagon is formed by N339 (Fig. 2c). Three of these five residues, N339, Y341, and Y443, are within the interdomain linkers. The carboxyl group of ET forms a hydrogen bond to the side chain of T296, and electrostatic interactions with K242 and R379 (Fig. 2d).

The C2-sulfur-imidazole moiety of ET protrudes from the pentagonal cage toward the opening of the ET binding cleft, with the imidazole ring aligned roughly parallel to the domain interface. Here, specific side chain-ligand interactions are relatively few, with E375 in the middle domain engaging N^ε2^ in a hydrogen bonding interaction, and P401 making van der Waals contact with the thione S (Fig. 2e). On the terminal domain, I243, the methyl group of T274, and the aliphatic part of the K242 side chain make van der Waals contacts with ET, while the hydroxyl group of T275 is close to N^δ1^ (Fig. 2f). Except for T274, these residues are largely conserved among EgtU sequences (see below).

A string of highly ordered, high occupancy water molecules essentially surrounds the thione-imidazole ring, making close contacts with the thione S and imidazole N^ε2^, while also bridging conserved tyrosines Y419 and Y341 that form two sides of the cage and bridge the middle and terminal domains (Fig. 2g). A network of nine high occupancy water molecules (Supplementary Table 2) extends from the side chain of E246 to the hydroxyl group of Y419 (Fig. 2e-g). This network is directly connected to a number of surface waters positioned in the cleft between the two domains which would provide an entropic driving force for the release of ET once docked to the permease domain dimer at the membrane, and further suggests that these waters might be considered co-ligands^67^. G244 is near the terminus of this extended network of water molecules (Fig. 2f) and is invariant in EgtU SBDs (see below). In the glycine betaine binding SBP ProX from *A. fulgidis*, G244 is replaced with phenylalanine, which would severely disrupt this network of largely buried water molecules (Fig. 2h). Furthermore, all *Af*ProX-family glycine betaine-specific SBPs use four Tyr to create the pentagonal cage, whereas all EgtUs have F293 in place of *Af*ProX Y63, which is accompanied by a switch of Y337 for F107 in *Af*ProX (Fig 2j). Substitutions here might be expected to negatively impact ET binding affinity, some via by disruption of this network of bound water molecules.

### Bound water molecules contribute to ET binding enthalpy

In some SBPs, water molecules lining the binding pocket have been proposed to contribute to ligand promiscuity^68^, while in others^69^, a highly ordered network of water molecules that form hydrogen bonds within a ligand binding pocket have been shown to contribute to a substantial enthalpic driving force for binding while imparting a high degree of selectivity. We therefore used isothermal titration calorimetry (ITC) to explore the thermodynamics of ET binding to EgtU SBD (Fig. 3a) and to assess the impact of mutations in perturbation of the network of bound water molecules. The equilibrium binding affinity (*K*_a_) of 1.7×10^7^ M^-1^ is consistent with the affinity obtained by tyrosine fluorescence. ET binding is enthalpically driven, with a Δ*H* comparable to Δ*G*, and a *T*Δ*S* value very close to zero (Fig. 3a, Table 1). These thermodynamic parameters are similar to those previously found for a taurine-specific SBP^70^, with an enthalpic driving force consistent with trimethylamine cation-π interactions found in osmoprotectant SBP-ligand complexes^60,62^. A G244F substitution mutant of EgtU binds ET ≈100-fold more weakly than wild-type EgtU SBD, with strongly perturbed thermodynamics (Fig. 3b); consistent with an important role of these water molecules in organizing the ET binding site, a finding previously observed in some other QAC-specific SBPs^67,70^. Two other substitutions that target conserved residues Y419 and F293 give rise to somewhat smaller perturbations of the binding energetics and affinity. The hydroxyl of Y419 is quite close to N^δ1^ of ET and might participate in hydrogen bonding. Indeed, mutation of Y419 to F causes a large reduction in the favorable binding enthalpy that is only partially compensated by favorable increase in the binding entropy, resulting in a ≈10-fold reduction in ET binding affinity (Fig. 3c). A F293Y *Sp*EgtU SBD substitution also reduces the affinity for ET by ≈10-fold, largely in the enthalpy term (Fig. 3d).

**Figure 3.**
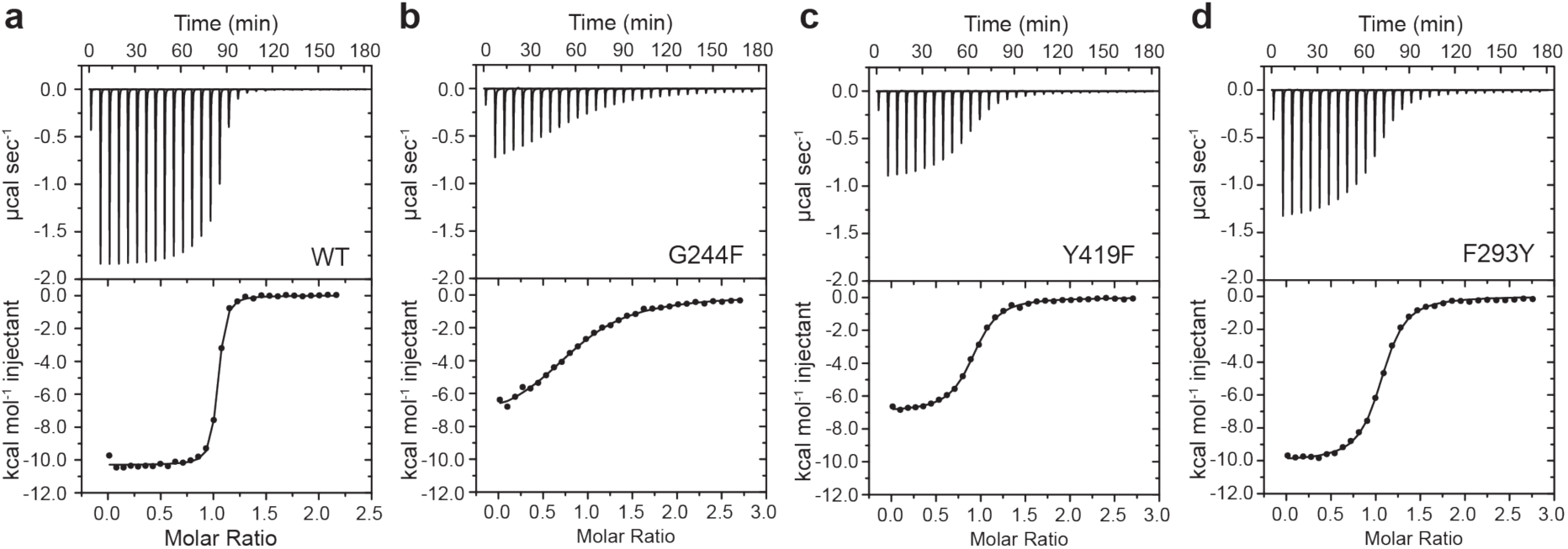
Thermodynamics of ET binding to wild-type and mutant *Sp*EgtU SBDs. **a-d**, Representative isothermal titration calorimetry-based titrations of WT and indicated mutant EgtU SBDs with ET, with thermodynamic parameters compiled in Table 1. Upper panel: Raw titration data. Lower panel: Dilution heat-corrected and concentration-normalized integrated raw data with the continuous line a best fit to a single-site binding model.

### ET induces a significant conformational change in the EgtU SBD

Although we were unable to obtain a structure of the ligand-free apo EgtU SBD, a structure of the *Sp*EgtU homolog *Listeria monocytogenes* BilEB involved in bile acid resistance (see below) is available in the apo state (PDB 4Z7E)^62^ and shows a much more open conformation. Our AlphaFold2^71^-generated model of apo-EgtU SBD shows a very similar “open” conformation (Fig. 4a) relative to the “closed” ligand-bound state. The middle domain and the terminal domain of the apo model each individually align closely with the corresponding domain in the ET-bound structure. Differences between the phi and psi backbone dihedral angles of the apo model and the ET-bound EgtU-CTT structure are limited and largely localized to a few loops and the linkers that connect the terminal and middle domains (Supplementary Figure 6a). However, these limited changes in the linkers are sufficient such that structural alignment of the terminal domains results in a displacement and a 54° rotation of the middle domain of the apo model relative to the same domain in the ET-bound structure. To validate the model and to assess whether ET drives an open-to-closed transition in solution, we prepared a F277W/L374C double mutant of EgtU SBD and attached a bimane group to C374 in the middle domain; here, the nonnative Trp in the terminal domain is expected to quench the bimane fluorescence to an extent dictated by the distance separating the two fluorophores (Fig. 4a-b)^72^. Titration with ET results in a significant quenching of the bimane fluorescence (Fig. 4c), consistent with ET-mediated closure driven by a rigid-body rotation/translation of one domain relative to the other (Fig. 4b). Moreover, the binding affinity of this construct is identical to that of wild-type EgtU SBD (Fig. 4d; Table 1).

**Figure 4.**
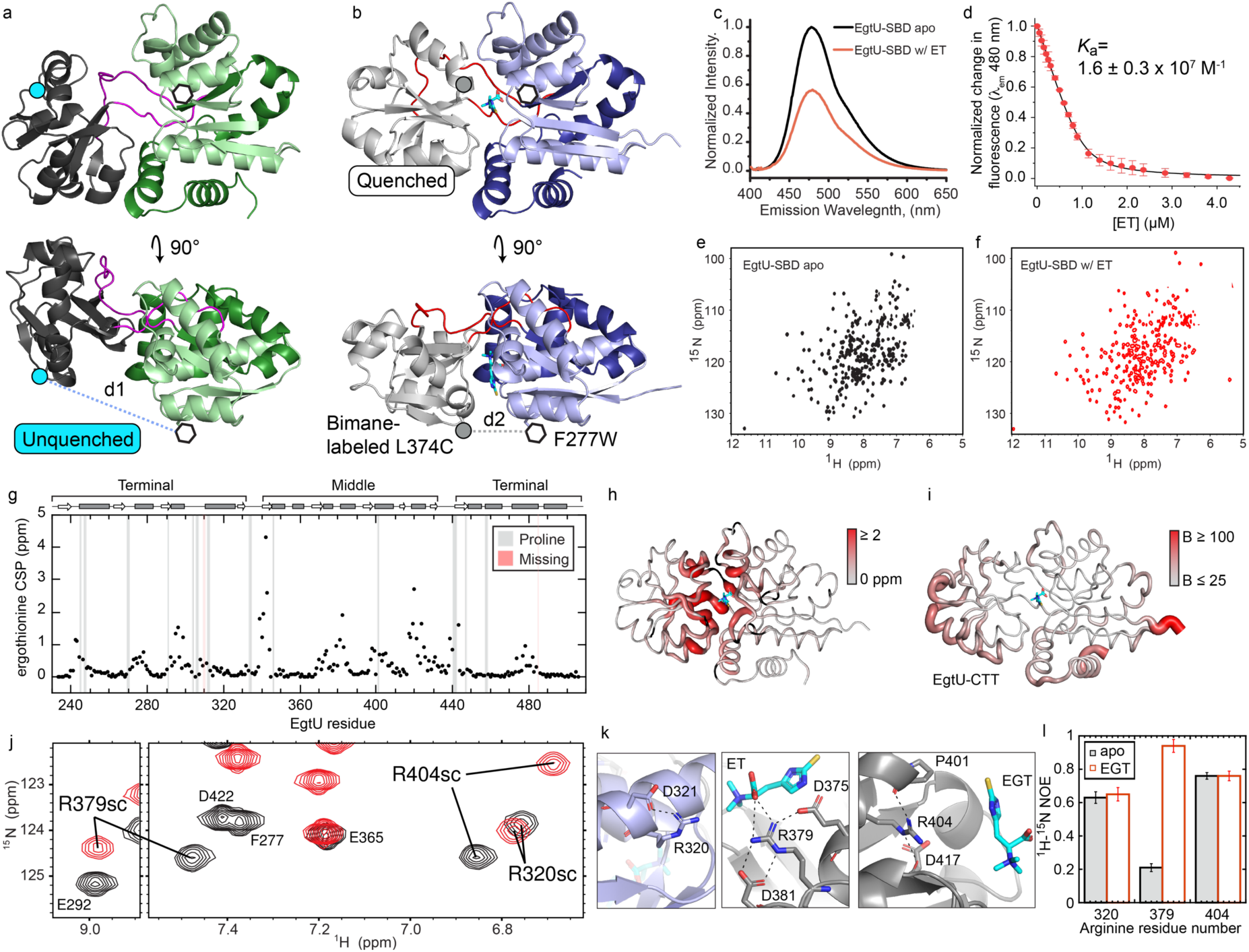
Conformational and dynamic changes in *S*pEgtU SBD upon ET binding. **a**, AlphaFold2 model of “open” apo *Sp*EgtU SBD (dark gray middle domain, green terminal domain, magenta linkers). The approximate site of the F277W substitution is indicated by an open hexagon, while the L374C site of the bimane labeling is shown as a cyan circle on the model. **b**, The “closed” crystal structure of ET-bound SBD (light gray middle domain, blue terminal domain, red linkers), with the F277W and L374C site of the bimane labeling shown as in panel **a. c**, Bimane fluorescence emission spectrum of bimane-labeled L374C/F277W *Sp*EgtU SBD with (red) and without (black) ET normalized the ligand-free spectrum. **d**, Normalized change in fluorescence emission at 480 nm upon titration of bimane-labeled L374C/F277W *Sp*EgtU SBD with ET with residue-specific assignments provided. **e**, ^1^H,^15^N TROSY spectrum of apo EgtU-SBD (Supplementary Figure 7). **f**, ^1^H,^15^N TROSY spectrum of EgtU-SBD bound to equimolar ET (Supplementary Figure 8). **g**, Backbone chemical shift perturbations (CSPs) upon binding ET for each residue in *Sp*EgtU SBD. Assignments are missing for residues H310 and V485 in the ET-bound state (shaded pink) with prolines shaded gray. **h**, CSPs of ET binding painted onto the crystal structure of ET-bound *Sp*EgtU SBD, with large chemical shift changes shown as thick, red tubes. **i**, B-factors plotted on the crystal structure of *Sp*EgtU SBD_CTT_, with high values shown as thick, red tubes, revealing low B-factors in the interdomain linkers, and comparable to the B-factors of high occupancy solvent molecules. **j**, Overlay of ^1^H,^15^N TROSY spectra of apo and ET-bound EgtU, zoomed in on the region where the arginine side chain peaks are folded into this spectral window. **k**, H-bond networks of arginine side chains (R320, R379, R404) found in the crystal structure. **l**, ^1^H,^15^N heteronuclear NOEs for these three arginine side chains.

### NMR studies of ligand-induced conformational change in the EgtU SBD

In order to understand the differences between the apo- and ligand-bound states of EgtU SBD in more detail, we turned to NMR spectroscopy. The 2D ^1^H,^15^N TROSY spectrum of apo EgtU SBD shows broad chemical shift dispersion, with uniform crosspeak intensities, consistent with a globular domain with an α/β fold (Fig. 4e). Addition of equimolar ET causes significant changes in the spectrum (Fig. 4f). Backbone chemical shift assignments of the apo- and ET-bound states of ^15^N, ^13^C, ^2^H-*Sp*EgtU SBD (Supplementary Figures 7-8) give rise to chemical shift-based secondary structure predictions of the ET-bound state that are identical to the crystal structure, and are strikingly similar in the apo state, consistent with our structural model that shows minimal changes in the individual subdomains. Closer inspection reveals that the apo state has an extended β–strand at the end of the first linker and in a neighboring strand in the middle domain (Supplementary Figure 6b). This small conformational change in the linker is consistent with the difference between the ET-bound structure and apo model, and is evidently sufficient to drive the opening and closing of the entire SBD. Chemical shift perturbations (CSPs) caused by equimolar ET binding are dramatic, but largely localized to loops in the domain interface, with the greatest changes in the linker and loop where conformational changes were identified (Fig. 4g-h).

The canonical “Venus fly trap” model for SBD substrate binding predicts that the apo protein is flexible, with the two subdomains rotating about the hinge, sampling multiple “open” conformations. Substrate binding then brings the two subdomains together, stabilizing a single “closed” conformation. With 10-residue linker regions and many interdomain contacts, one might expect the linker to display some flexibility even in the bound state. However, crystallographic B-factors are low (Fig. 4i) and ^15^N[^1^H] heteronuclear nuclear Overhauser enhancements (hNOE) are high throughout the linkers in the bound state, indicating that the linker is strikingly rigid when bound to ET. Moreover, high hNOEs suggest that the linker is also rigid in the apo state (Supplementary Figure 6c); in general, binding of ET to EgtU has strikingly little impact on sub-ns backbone dynamics throughout the molecule (Supplementary Figure 6c-d).

In order to compare the backbone dynamics of the apo and ET-bound states of EgtU SBD in more detail, we assessed several structural models of the apo state. At one extreme, we used a model identical to the ET-bound crystal structure. The model at the other extreme holds that the middle and terminal domains tumble independently of one another, connected by an infinitely flexible linker. The AlphaFold2 model represents an intermediate condition, structurally distinct from the ET-bound state, yet completely rigid. We used HYDRONMR to predict the ^15^N *R*_1_ and *R*_2_ backbone relaxation parameters for each model and compared these ratios to the corresponding ratio of experimental parameters. Overall, the fitted experimental values show relatively slow tumbling for the molecular weight (Supplementary Figure 6d). As expected, the predictions based on the ET-bound crystal structure match best to the experimental parameters for ET-bound EgtU SBD in solution (Supplementary Figure 6e) relative to the experimental parameters for apo EgtU SBD (Supplementary Figure 6f).The apo experimental data match significantly better to the fully rigid AlphaFold2 model (Supplementary Figure 6g) than to either the ET-bound crystal structure or the model with fully uncoupled domains (Supplementary Figure 6h). These data reveal that ET binding triggers a rigid-body transition from a conformationally narrow open state to another conformationally narrow closed state, in striking contrast to a conformational selection model where the apo-state ensemble is heterogeneous.

Although the backbone relaxation parameters in EgtU SBD (Supplementary Figure 6i-j) are strikingly insensitive to ET binding, side chains in the binding pocket are quite sensitive to ligand binding. Arginine guanidino protons are rarely observable in ^15^N,^1^H-TROSY spectra acquired at pH 7.0 due to their rate of solvent exchange. Surprisingly, three slowly exchanging guanidino protons are observable (Fig. 4j), with hydrogen bonds in the crystal structure consistent with a slowed exchange rate (Fig. 4k). R320 forms H-bonds with D321 far from ligand binding site, while R404 H-bonds with P401 found in the cleft near the bound ET. R379 is sandwiched between E375 (conserved as E/D; see below) and the invariant D381 side chains, and forms a H-bond to the carboxylate oxygen of ET. The hNOEs of the side chains of R320 and R404 are unaffected by ET binding, but the hNOE of R379 is low in the absence of ligand, consistent with some motional disorder, and becomes strongly rigidified on the sub-ns timescale upon binding ET (Fig. 4l).

### The EgtU SBD binding of ET is highly specific

To explore the ligand specificity of EgtU SBD, we created a EgtU SBD-sfGFP fusion protein (Fig. 5a), the fluorescence of which is strongly quenched upon ET binding, and which binds ET with an affinity similar to wild-type SBD (Fig. 5b-c; Table 1). Using the same assay, we find that *L*-hercynine binds ≈10,000-fold less tightly (Fig. 5d). We then used this assay to screen the ability of other potential ligands (Supplementary Figure 9a) to quench the fluorescence of the EgtU SBD-sfGFP fusion protein and find that none do so, at concentrations 1000-fold higher than the *K*_d_ for ET (Fig. 5e), nor do they negatively impact the ability of ET to quench the fluorescence of EgtU SBD-sfGFP at 100-fold molar excess ligand relative to ET (Fig. 5f). ITC confirms results obtained by fluorescence spectroscopy, with no detectable change in heat observed for selected other ligands, even for the structurally closest mimic *L*-hercynine which binds weakly despite lacking only the thione sulfur atom of ET (Supplementary Figure 9b).

**Figure 5.**
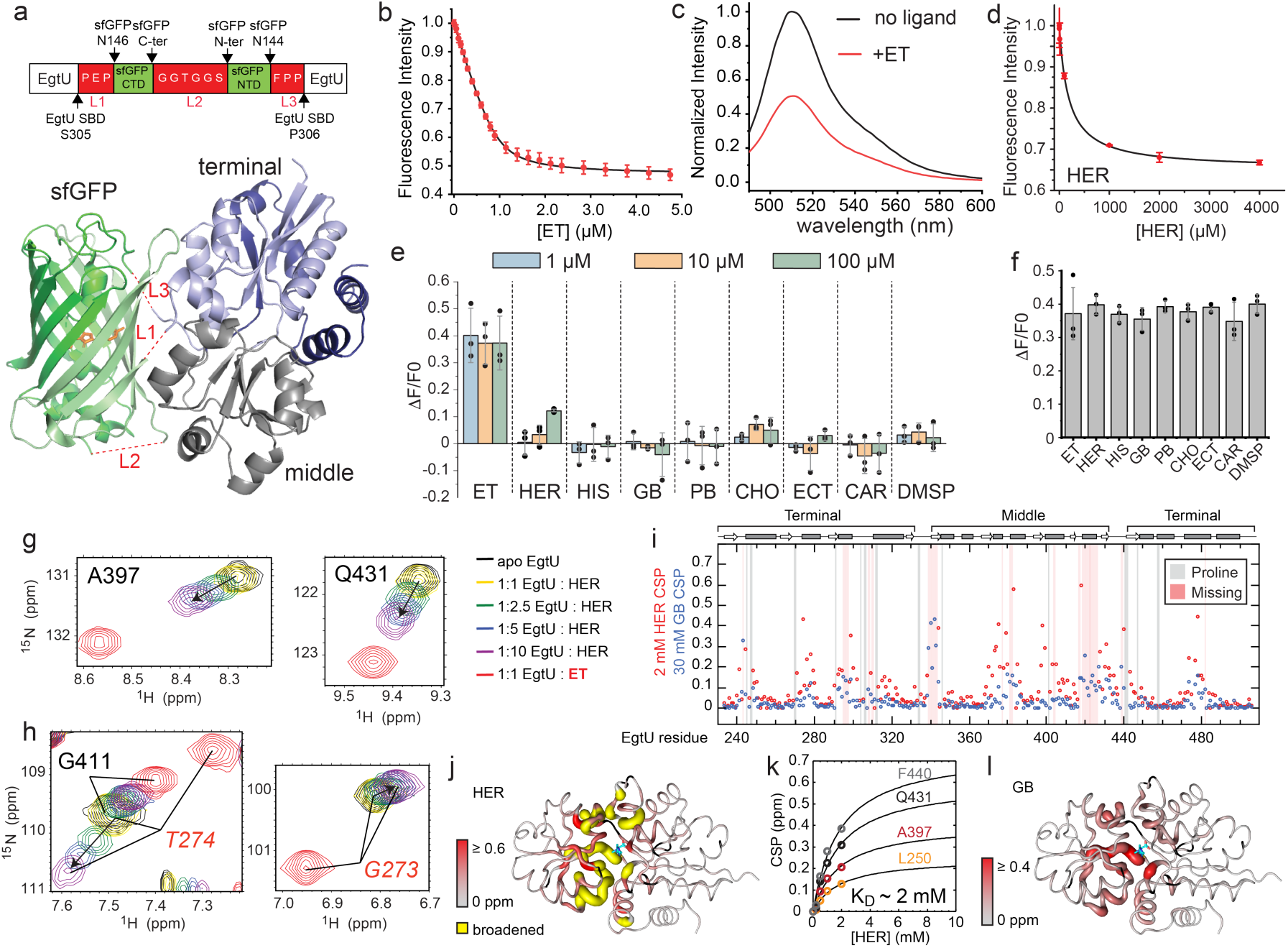
The *Sp*EgtU SBD binds ET with high selectivity. **a**, *Upper*, schematic representation of the EgtU SBD-sfGFP fusion protein construct; *lower*, Swiss model-generated cartoon, with the middle and terminal domains of EgtU SBD and loops L1-L3 indicated. **b**, ET binding to the EgtU SBD-sfGFP fusion protein, monitored by quenching of the sfGFP fluorescence. Continuous curve, fit to a 1:1 binding model. **c**, sfGFP fluorescence emission spectra, /∴_ex_=470 nm with or without 10 µM ET. **d**, same as (c) except *L*-hercynine was added to EgtU SBD-sfGFP fusion protein (Table 1). **e**, Quenching of sfGFP fluorescence of the EgtU SBD-sfGFP fusion protein following addition of 1, 10 or 100 µM of the indicated ligand. Each bar represents triplicate measurements with each data point represented by a filled circle. **f**, Same as (e), except that 1 µM ET (left bar) was compared to a mixture of 1 µM ET and 100 µM of the indicated ligand (other bars). Each bar represents triplicate measurements (filled circles). **g** and **h**, Movement of the indicated backbone NH crosspeak from the apo-state (black) as *L*-hercynine (HER) is added (yellow to purple), compared to the crosspeak position of ET-bound EgtU SBD (red). **i**, Backbone chemical shift perturbation (CSP) maps resulting from the addition of 2 mM HER or 30 mM glycine-betaine (GB). **j**, Backbone CSP maps upon HER binding painted onto the *S*pEgtU SBD structure. k, Fits to a 1:1 binding model for selected NH cross-peaks as a function of [HER] and globally fitted to a 1:1 binding model. l, Backbone CSP maps upon HER binding painted onto the *S*pEgtU SBD structure.

NMR was next used to probe the binding of weakly binding ligands to EgtU SBD in more detail. A titration of *L*-hercynine into ^15^N-labeled EgtU SBD shows that the ligand-bound and free conformations are in fast-to-intermediate chemical exchange on the ^1^H NMR timescale, with most peaks generally moving towards the corresponding resonance frequency of the ET-bound residue (Fig. 5g), while many vanish entirely. Only a few resonances, *e*.*g*., G273 and T274, shift in a direction that is opposite to ET (Fig. 5h); these residues are in close proximity to the thione sulfur atom (Fig. 2f). A large molar excess of ligand shows clear evidence of specific binding, with CSPs localized to the same interfacial loops that respond to ET (Fig. 5i,j; Supplementary Figure 10a), but 2 mM hercynine was insufficient to saturate EgtU SBD. Fitting the chemical shift perturbations for several residues as a function of ligand concentration gives an affinity estimate of 600 M^-1^ (Table 1). Titration of glycine-betaine shows only fast chemical exchange, consistent with even weaker binding, as even 30 mM glycine betaine fails to reach saturation (Supplementary Figure 10b). Largely the same binding pocket residues are affected (Fig. 5i,l), with an affinity estimated to be less than 30 M^-1^ (Table 1).

### EgtUV homologs are widely distributed across the genomes of firmicutes

We next asked if EgtUs cluster in a global sequence analysis, while also elucidating conserved features of an EgtU and how this differs from other osmoprotectant transporters. To do this, we used SPD_1642 as query to construct a sequence similarity network (SSN) using genomic enzymology tools^73,74^ to visualize the relationships among EgtU homologs in the context of the entire superfamily of osmoprotectant uptake (*opu*) SBPs/SBDs (Supplementary Figure 11). We find that EgtU is representative of a distinct subcluster within cluster 2 (Fig. 6a) containing sequences within individual metanodes that are characterized by the largest neighborhood connectivity of the entire SSN map (Fig. 6b; Supplementary Figure 12). Remarkably, *Sp*EgtU homologs are found nearly exclusively in firmicutes and include gastrointestinal tract-resident bacteria, notably *Lactococcus lactis*, and a wide range of human opportunistic pathogens beyond *S. pneumoniae*, including pathogenic *Bacillus* ssp., *B. cereus* and *B. infantalis* (previously OpuF^75^), *Enterococcus faecalis, Neisseria mucosa, Staphylococcus aureus* and *Listeria monocytogenes* (Supplementary Figure 13). A sequence logo representation of conserved residues (Fig. 6c) of the SBD reveals that all functional features described above in the *Sp*EgtU SBD-ET complex are highly conserved or invariant. These include those residues that define the QAC pentagonal cage and associated “second sphere” interactions, and those residues that interact with the imidazole and thione sulfur moieties of ET. On the other hand, EgtU homologs found in the central SSN cluster 2 subcluster, exemplified by *Clostridioides difficile* OpuF (Supplementary Figure 14),^76^ appear to harbor the full collection of mutations in *Sp*EgtU that individually reduce ET affinity. As a result, OpuF may be specific for another QAC, or simply exhibit relaxed QAC specificity. The ligand specificity of the other subclusters is not yet known.

**Figure 6.**
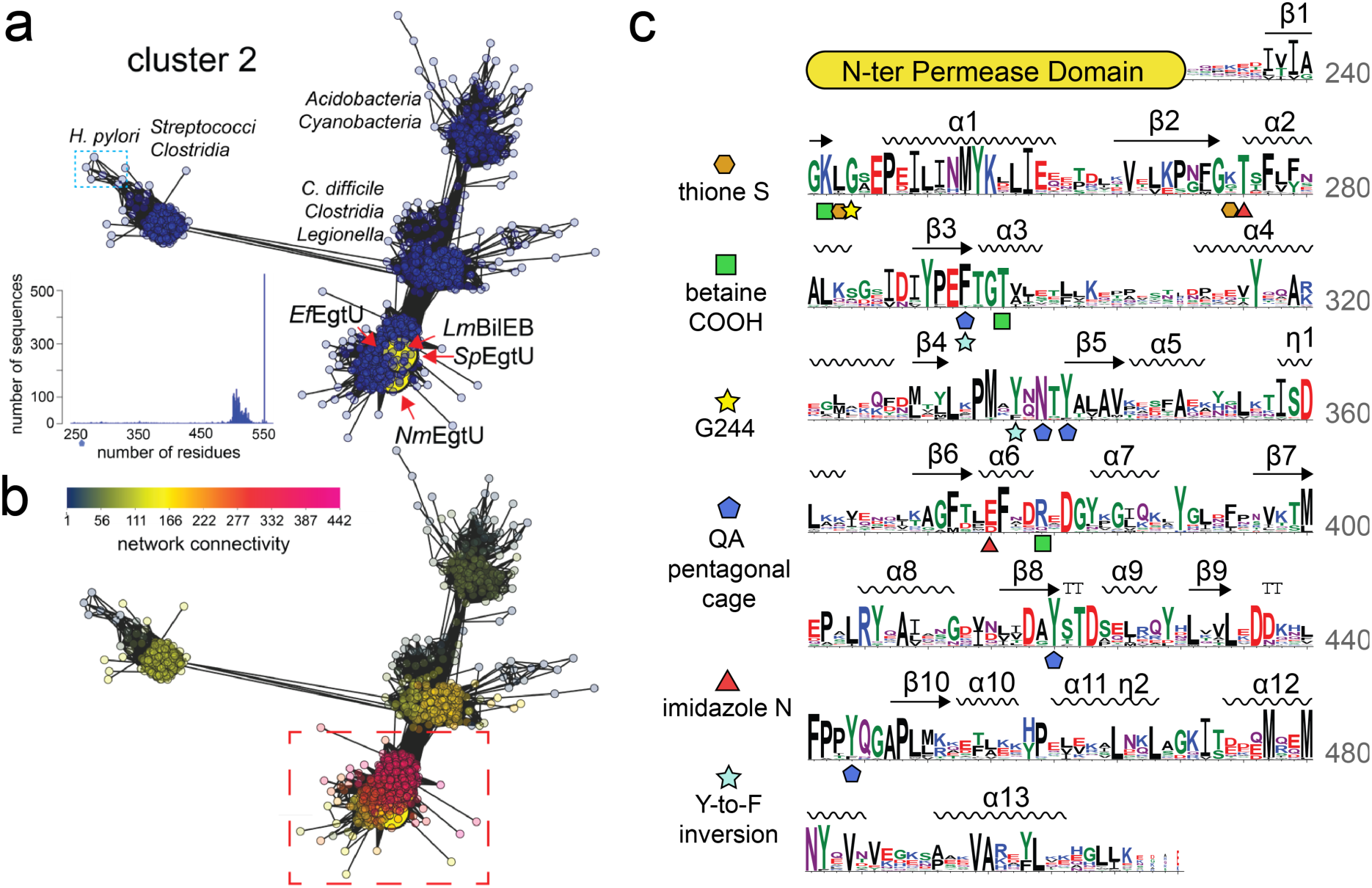
EgtU homologs cluster in a grouping of highly similar sequences within a subcluster of SSN cluster 2. **a**, Several EgtU sequence metanodes are highlighted with a large yellow circle, including *L. monocytogenes* BilEB and *E. faecalis* EgtU SBD biochemically characterized here. The middle subcluster sequences are derived from anaerobes or obligate anaerobes, including those recently studied in *C. difficile*^76^. *Inset*, length distribution of sequences within all of cluster 2. **b**, Representation of the SSN network colored according to neighborhood connectivity (NC; see scale), with those sequences within a single metanode that are most closely related characterized by a large NC index (and shaded *magenta*). **c**, Sequence logo representation of conservation in SBD of EgtUs of those closely related sequences encircled by the *red* box in panel **b**. Secondary structure of the SBD is indicated, based on the structure of *Sp*EgtU SBD. Residues discussed are highlighted with a specific symbol.

As a direct test of this functional grouping of proposed ET transporters, we purified and characterized the candidate EgtU SBD from *E. faecalis, S. aureus* and *L. monocytogenes*. We find that *Ef*EgtU SBD binds ET with an affinity comparable to that of *Sp*EgtU SBD (Supplementary Figure 15a; Table 1), while NMR spectra of apo- and ET-bound *Ef*EgtU SBD show similar features that are broadly consistent with comparable conformational changes to those described for *Sp*EgtU (Supplementary Figure 15b-c). Since ET is obtained in the diet in animals, these findings with *E. faecalis* EgtU might suggest a competition for ET among resident microbiota and opportunistic pathogens in the GI tract for some physiological advantage. Similar experiments were carried out with *Sa*EgtU SBD and we find a similar binding affinity, and no detectable binding by ITC to *L*-hercynine (Table 1; Supplementary Figure 15d). The EgtU homolog from *L. monocytogenes*, denoted BilEB, has long been known to be associated with bile acid resistance^56^ and early work ruled out a role for BilEB in the uptake of choline, carnitine or glycine-betaine^62^. We show here that *Lm*BilEB binds ET with an affinity identical to that of authentic *Sp*EgtU SBD (Table 1; Supplementary Figure 15e), which argues strongly that BilEB is ET uptake transporter.

## DISCUSSION

In this work, we show that *Sp*EgtUV possesses characteristics of a bacterial ABC transporter that is specific for the low molecular weight thiol/thione ergothioneine (ET). We show that deletion of *S. pneumoniae spd_1642* creates a strain that is unable to accumulate ET either when grown in a vertebrate tissue-derived growth media that contains significant endogenous ET, or on a chemically defined medium to which ET has been added. Our studies reveal that the solute binding domain of EgtU exhibits high selectivity for ET over even closely related QACs, *e*.*g*., *L*-hercynine, and is consistent with our functional assignment, supported by biochemical experiments of EgtU homologs from three other firmicutes. Detailed NMR experiments show that non-cognate QACs simply fail to stably close the ligand binding cleft between the two domains, a remarkable finding given that hercynine differs from ergothioneine only by the loss of the thione S. Given a paucity of direct contacts between *Sp*EgtU SBD and the thione S atom, we hypothesize that the extensive network of ordered water molecules must play an important role in ET complex formation, consistent with our characterization of G244F *S*pEgtU SBD (Table 1).

A comparative sequence analysis suggests that EgtU is widely distributed among firmicutes known to colonize the vertebrate gastrointestinal (GI) tract, including commensals and pathogens, as well other pathogens known to infect other tissues but also capable of replicating in immune cells. Indeed, a recent report shows that the gut commensal bacterium, *Lactobacillus reuteri*, readily takes up extracellular ET, although the mechanism of uptake was not defined in that study^77^. It is well established that commensals resist colonization by pathogens in the gut by depleting essential nutrients and remodeling resource allocation in this niche^78^; the work reported here raises the strong possibility of a competition between commensal and pathogen for a nutrient that may well be protective against oxidative and antibiotic stressors.

Our findings clearly establish a mechanism by which a bacterium need not synthesize ET in order to access its potential antioxidant properties^36,79^. In this case, ET is likely obtained in the diet of the vertebrate host where GI-resident bacteria that express EgtUV would have initial access to this metabolite. As described above, the human ET transporter (ETT) is expressed in a wide range of tissues and cells, including neutrophils and macrophages^48^; this suggests that ET may be bioavailable to both extracellular and intracellular pathogens, *e*.*g*., those phagocytosed by neutrophils. This is not yet known with certainty since concentrations of ET itself have not been comprehensively mapped in a wide variety of tissues or cells in an infected host using the analytical approaches we describe here. However, bacteria found in either intracellular or extracellular lifestyle may well be capable of capturing ET, given that ETT is reported to bind ET with a *K*_d_ of 50 µM, which is ≈1000-fold weaker than the bacterial EgtU SBD described here.

What role ET plays in bacterial cell physiology can be hypothesized from literature published prior to the knowledge that EgtUV homologs encode an ET transporter. For example, deletion of *egtUV* gives rise to a fitness defect in a lung infection model in *S. pneumoniae* D39 strain and is thus a virulence factor^80^. We provide evidence to suggest that ET is the long-sought metabolite that is transported by *Listeria monocytogenes* BilEB, required to promote an adaptive response to bile acid stress during gastrointestinal transit^56^. How ET protects *L. monocytogenes* from bile acid stress is unknown but oxidative stress resistance is a strong possibility. In methicillin-resistant *S. aureus, Sa*EgtUV expression is induced ≥15-fold after long exposures to human neutrophil-derived azurophilic granule proteins, but with no significant response to peroxide and hypochlorous acid stress at the same time points^81^; this suggests an as yet unknown ET-dependent phagocytosis resistance mechanism to killing by these effectors. In *E. faecalis, egtUV* are among the most highly upregulated genes in bacteria isolated from a simplified human microbial consortium (SIHUMI) colonized IL10 (interleukin-10)^−/−^ mice vs. wild-type mice. This increase in expression is lost when *E. faecalis* is monocolonized in IL10^−/−^ mice, a finding that suggests competition for this thiol may be physiologically important^82^. Finally, although recent studies show that EgtU (denoted OpuF) from *Bacillus infantis* and *Bacillus panaciterra*, is capable of rescuing an osmoprotectant uptake-deficient *B. subtilis* strain grown in hyperosmotic conditions, the concentrations required to do this are in the high μM to mM range on a chemically defined growth medium^75^. This finding is consistent with the very weak, but measurable (*K*_d_≈mM) binding of EgtU SBD to glycine betaine and *L*-hercynine observed here. On the other hand, we have not yet explored the potential impact of oxidation or methylation of the ET thione S to create sulfonylated ET, 5-oxo-ET or *S*-methylated ET, respectively^83^, or Se substitution of the S atom in ET^84^, on EgtU SBD ligand binding affinity. Indeed, the ligand specificity of EgtU homologs in other SSN cluster 2 SBDs remains to be experimentally validated^76^.

Beyond the function of ET itself, the mechanism by which EgtUV is upregulated may also provide insights in the pathogen response to host effectors, especially in oxidative stress adaption. In *M. tuberculosis*, the biosynthesis of ET is regulated by the ROS- and RNS-sensing 4Fe-4S cluster transcriptional regulator WhiB3^40^, with the bacterial concentration of ET increasing ≈7 fold in a Δ*whiB3* strain^79^. In addition, the ET level is significantly increased in Δ*whiB3* when fatty acids serve as the nutritional carbon source^39^. Although ET is present at lower concentrations than the major LMW thiol in *M. tuberculosis*, mycothiol, ET becomes significantly elevated in a mycothiol-biosynthesis deficient strain. Our LMW thiol profiling also confirms that ET is present at a significantly lower cellular level relative to glutathione in *S. pneumoniae*, but distinct from other organisms, *S. pneumoniae* is totally dependent on scavenging LMW thiols from its immediate microenvironment to meet cellular needs. How the pneumococcus and other firmicutes balance EgtUV-mediated uptake of ET vs. other thiols is unknown. How the pneumococcus regulates *egtUV* expression is also not yet known, although an uncharacterized dithiol-containing MarR (SPD_1645) is found in the operon harboring *egtUV* (Fig. 1a) whose expression is clearly tied in some way to quinone-derived oxidative stress, mediated in part by catechol-Fe^III^ uptake^57^. The known ability of ET to form coordination complexes with Fe^II^, may suggest a secondary role in suppressing redox cycling of iron by host-derived H_2_O_2_ and other potent ROS and RNS or some other role in colonization^27^ or virulence.

## METHODS

### Reagents

*L*-glutathione (GSH) was obtained from Sigma Aldrich (G4251), *L*-ergothioneine from Santa Cruz Biotechnology (sc-200814), *L*-cysteine from Fisher Biotec (BP376-100), *L*-hercynine from Toronto Research Chemicals (H288900), *L*-histidine from Sigma Aldrich (H8000), choline chloride from Sigma Aldrich (C1879), glycine-betaine from Sigma Aldrich (B3501), ectoine from Sigma Aldrich (81619), proline-betaine from VWR (TCS0358), *L*-carnitine from Sigma Aldrich (C0283), and dimethypropiothetin hydrochloride from Sigma Aldrich (80828). These chemicals were used without further purification. Other chemicals include IPTG from GoldBio (2481C100), TCEP from Chem-Impex (00194), dithiothreitol from Chem-Impex (00127), Tris-HCl from MP Biomedicals (816100), HEPES from Chem-Impex (00174), EDTA from VWR Chemicals (BDH9232), NaCl from VWR Chemicals (BDH9286), Imidazole from Chem-Impex (00418) and monobromobimane (mBBr) from Sigma (B4380). BHI broth was obtained from BD (37500, lot 1159859) while Luria broth was obtained from Fisher Bioreagent (BP9723). Milli-Q water was used to make all solutions.

### *Streptococcus pneumoniae* D39 mutant strain preparation and growth conditions

The mutant strains listed in Supplementary Table 1 were constructed using standard laboratory practices for allelic replacement in *S. pneumoniae* serotype 2 D39W (IU1781)^85^. All mutant strain constructs were sequence verified. Primers are listed in Supplementary Table 4.

### LMW thiol profiling in *Streptococcus pneumoniae* D39

Bacterial cell pellets for LMW thiol profiling were prepared by inoculating selected strains in BHI medium under microaerophilic conditions with 5% CO_2_ at 37 °C as described in our previous work^86^. In a chemically defined medium (CDM)^87,88^, cells were grown in triplicate in the same condition with or without addition of ET to the indicated concentration (0, 0.05, 0.5, 5 μM). Cells from 5 mL culture were collected at OD_620_ ∼0.2-0.3. Cell pellets from 4 mL of this culture were extensively washed in chilled PBS and frozen in –80°C for LMW thiol profiling. Cell pellets from the remaining 1 mL were washed with chilled PBS and immediately frozen at –80 °C for cellular protein quantification.

Heavy (*d*_4_) HPE-IAM was used to create alkylated selected LMW thiol standards^89^. The cellular LMW thiols were alkylated by light (H_4_) HPE-IAM and quantified by LC-MS by spiking a known concentration of LMW thiol derivatized by heavy *d*_4_-HPE-IAM as a standard. *d*_4_-HPE-IAM and HPE-IAM were chemically synthesized as described and structural integrity confirmed by NMR spectroscopy^89^. In brief, each cell pellet was resuspended in 100 µL Milli-Q water with 5 mM HPE-IAM by adding 1 μL 0.5 M HPE-IAM stock prepared in DMSO. The resuspended cell pellet was lysed using a 1 min freeze in liquid N_2_ and 37 °C water bath thaw for 1 min. Five freeze-thaw cycles were performed and cell lysates and further incubated at 37 °C for 1 h, and then microcentrifuged at top speed for 20 min. 50 μL supernatants were transferred and filtered by 0.2 μm cutoff micro-centrifuge filter tubes. Then 1 µM heavy *d*_4_-HPE-IAM-derivatized LMW thiol standards (ET, cysteine, GSH) were added into the flow through with total volume brought up to 100 µL with Milli-Q water. Both light and heavy *d*_4_-HPE-IAM labeled standards were prepared by capping 100 μM reduced LMW thiols (ET, cysteine, GSH) with 3 mM *d*_*4*_-HPE-IAM in the lysis buffer at 37 °C for 1 h (see Supplementary Figure 1). The samples were analyzed by a C18 (YMC-Triart C18) LC system coupled to a Waters SYNAPT G2S high resolution MS using a mobile phase A (5% acetic acid, 10% methanol) and mobile phase B (5% acetic acid, 90% methanol) with the following LC elution gradient: 0-3 min, 100% A, 0% B; 3-7 min, linear gradient to 75% A, 25% B; 7-9 min, 75% A, 25% B; 9-12 min, linear gradient to 25% A, 75%B; 12-14 min, linear gradient to 0% A, 100% B; 14-20 min, 0% A, 100% B. The resulting total ion chromatogram (TIC) was searched for positively charged ions (*z*=1; M^+^ or M+H^+^) (mass tolerance of ±0.02 *m/z*) and the extracted ion chromatograms of each light (H_4_) and heavy (*d*_4_) HPE-IAM-capped thiol identified in MS1 obtained, peak areas quantified, and identity confirmed by LC-MS/MS by comparison to the corresponding authentic compound standard (Supplementary Figures 2-4). The ratio of the light and heavy MS1 features was used to calculate the concentration of each thiol using the known concentration heavy standard spiked into the mixture. The remaining 1 mL culture cell pellets were analyzed by Bradford Assay to quantify the total protein concentration of each sample as described in previous work^90^. The LMW thiol concentration is presented as nmol/mg total protein.

### Cloning, protein expression, and purification of EgtU SBDs from *S. pneumoniae, Enterococcus faecalis, Staphylococcus aureus* and *Listeria monocytogenes*

The regions of the genes encoding the soluble, extracellular C-terminal solute binding domain (SBD) of *S. pneumoniae* D39 EgtU (locus tag SPD_1642) from residue E233 was PCR-amplified from the genomic DNA. The same was done for the candidate EgtU SBDs of *E. faecalis* OG1RF EgtU (locus tag RS02210) beginning at residue K233, *S. aureus* FPR3757 USA300 (locus tag *sau300_0707*) beginning at residue G233, and *L. monocytogenes* strain 10403S (locus tag *lmo1422*) beginning at residue S231. The primers used in the cloning are listed in Supplementary Table 4. Each gene was inserted into the pSUMO expression vector^91^ with an N-terminal hexa-histidine tag, with mutants prepared by PCR-based site-directed mutagenesis. The *Sp*EgtU-SBD-sfGFP expression construct was prepared using primers (Supplementary Table 4) largely following a published procedure^92^. In brief, the PCR fragment F1 containing *Sp*EgtU-SBD and pSUMO plasmid (6.5 kB) was prepared using *Sp*EgtU-SBD pSUMO expression vector as template and primers *Sp*EgtU_S_P1 and *Sp*EgtU_S_P2. PCR fragments containing CTD or NTD of sfGFP with linkers were prepared by using primer pairs *Sp*EgtU_S_P3/*Sp*EgtU_S_P4 and *Sp*EgtU_S_P5/*Sp*EgtU_S_P6. The genomic DNA of IU9985 containing sfGFP DNA sequence was used as a template^93^. The sfGFP CTD and NTD fragments were linked together by fusion PCR using primer pairs *Sp*EgtU_S_P3/*Sp*EgtU_S_P6 to generate fragment F2. Fragment F2 was ligated to fragment F1 by a Gibson assembly protocol^94^. Expression vectors were amplified in *E*.*coli* DH5α and sequences verified.

The sequence verified expression vectors were transformed into *E. coli* BL21(DE3) and grown in either LB (*S. pneumoniae, E. faecalis*) or an M9 minimal medium (*L. monocytogenes, S. aureus*) supplemented with 30 µg/mL kanamycin. 1 mM isopropyl β-*D*-1-thiogalactopyranoside (IPTG) was added to induce protein expression at OD_600_≈0.8. Following overnight expression at 18 °C, the cells were pelleted by centrifugation. The cell pellet was resuspended in Buffer A (25 mM Tris-HCl, pH 8), 500 mM NaCl, 10% glycerol, 20 mM imidazole) and lysed by sonication on ice. The crude lysate was clarified by centrifugation. 70% ammonium sulfate was applied to precipitate the protein and the pellet was collected by centrifugation. The precipitated pellet was resuspended in Buffer A and the solution subjected to Ni(II) immobilized affinity chromatography using a 5 mL HisTrap FF column (GE Healthcare Life Sciences) with a gradient from 100% buffer A to 100% buffer B (25 mM Tris-HCl, pH 8.0, 500 mM NaCl, 10% glycerol, 500 mM imidazole). The fractions containing the His-tagged SUMO fusion protein were pooled and digested by SUMO protease (20 µg/mL) while dialyzing in buffer A with 2 mM dithiothreitol (DTT) at room temperature. The digested protein fractions were applied to a HisTrap FF column in Buffer A. The flow-through fractions were pooled and concentrated by centrifugation with a 10 kDa cutoff and subjected to size exclusion chromatography on a Superdex-200 column in Buffer C (25 mM Tris-HCl, pH 8.0, 500 mM NaCl, 2 mM EDTA) and monomeric fractions pooled. The concentration of purified protein was measured using the molar extinction coefficients at 280 nm (ε_280_) (Supplementary Table 5). Purified protein fractions were pooled and stored at –80 °C until use.

### Intrinsic tyrosine fluorescence titration analysis

Data were acquired on a PC1 spectrofluorometer with λ_ex_ 285 nm (2 mm slit) and the emission intensity recorded through a 305 nm cut-off filter. The ligand was prepared in titration buffer (50 mM HEPES, pH 7.5, 150 mM NaCl, 2 mM EDTA). All proteins were buffer exchanged into same titration buffer and ligands were titrated into 3 mL 1 µM protein. The titrations were carried out with continuous stirring at 25.0 (±0.1) °C and resulting data corrected for dilution and the inner filter effect and fit to a 1:1 protein:ligand binding model to estimate *K*_*a*_ using DynaFit^95^.

### Isothermal calorimetry titration

ITC experiments were carried out using a MicroCal VP-ITC calorimeter at 25 (±0.1) ºC by titrating 20 or 30 µM *Sp*EgtU SBD or the indicated mutant in the sample chamber in 50 mM HEPES, pH 7.5, 150 mM NaCl, 2 mM EDTA with the indicated ligand (ET, *L*-hercynine, *L*-histidine or glycine-betaine) in the syringe in same buffer. For the ET titration, the ligand concentration in the syringe was typically 375 μM with 30 μM protein in the sample chamber. For other ligands, the ligand concentration was 600 μM, and 20 μM protein in chamber. The raw ITC data were integrated, concentration normalized, and plotted as heat versus metal/protein ratio using Origin. All data were fit to a single site binding model included in the data analysis package provided by MicroCal.

### Protein crystallography and data analysis

The purified protein was buffer exchanged into crystallography buffer, 50 mM Tris-HCl, 150 mM NaCl, 2 mM EDTA pH 7.5. A 3-fold excess of ergothioneine was added to the purified protein and excess ET removed by chromatography on a HiLoad 16/600 Superdex 200 size exclusion column (Cytiva). The main peak corresponding to monomeric *Sp*EgtU SBD was pooled and concentrated for protein crystallography screening. *Sp*EgtU SBD_CTT_–ET (15 mg/mL) crystals grew in sodium citrate, pH 5.6, 0.2 M potassium sodium tartrate and 1.8–2.0 M ammonium sulphate at 20 °C using the hanging-drop vapor-diffusion method. *Sp*EgtU SBD-ET (15 mg/ml) crystals grew in 1.6 M sodium citrate, pH 6.5, at 20 °C using the hanging-drop vapor-diffusion method. Crystals were harvested, cryo-protected in a reservoir solution supplemented with 25% glycerol and flash-frozen in liquid nitrogen. Diffraction data were collected at 100 K at the Beamline station 4.2.2 at the Advanced Light Source (Berkeley National Laboratory, CA) and were initially indexed, integrated, and scaled using XDS^96^. Molecular replacement was used to estimate phases using PHASER and PDB code 4Z7E^62^ as search model. Successive cycles of automatic building in Autobuild (PHENIX) and manual building in Coot, as well as refinement (PHENIX Refine) led to complete models^97^. MolProbity software^98^ was used to assess the geometric quality of the models, and Pymol was used to generate molecular images. Data collection and refinement statistics are indicated (Supplementary Table 1).

### Tryptophan-mediated quenching of bimane fluorescence

F277W/L374C *Sp*EgtU SBD was prepared as described above except that buffer for protein purification was degassed and 2 mM TCEP was added to all purification buffers. The purified protein was then buffer exchanged into degassed labeling buffer, 200 mM sodium phosphate, Ph 7.4 without reducing reagent. A mBBr stock solution was prepared in DMSO at 20 mM and stored at –80 ºC until used. 20 µM protein was mixed with a 30-fold molar excess of mBBr in labeling buffer at 37 °C for 1 h and excess mBBr removed by eight rounds of washing through a 10 kDa cut-off centrifugation filter with labeling buffer. The concentration of the labeled protein was measured by absorption at 280 nm using the molar extinction coefficient shown (Supplementary Table 5). The conjugated bimane concentration was measured using an ε_380_ of 5000 M^-1^ cm^-1^. Data were acquired in 50 mM HEPES, pH 7.5, 150 mM NaCl, 2 mM EDTA on a PC1 spectrofluorometer with λ_ex_ 380 nm (2 mm slit) and the emission intensity recorded through a 480 nm cut-off filter. Ligands were prepared in this buffer with the protein buffer-exchanged into the same buffer. Various ligands were titrated into 3 mL 1 µM protein up to 5 µM total ligand, with continuous stirring at 25.0 (±0.1) °C. All titration data were fit to a 1:1 protein:ligand binding model to estimate *K*_a_ using DynaFit^95^. The total emission spectrum was acquired from 400 nm to 650 nm before and after the titration, with the initial emission intensity at 480 nm normalized to 1 and that at 650 nm normalized to 0.

### NMR backbone assignments

Uniformly ^15^N, ^13^C, ^2^H -labeled protein was expressed in *E. coli* BL21 (DE3) cells in M9 minimal medium containing 1 kg D_2_O, as well as 1.0 g of ^15^NH_4_Cl and 2 g ^13^C_6_,^2^H-glucose as the sole nitrogen and carbon sources, respectively. Uniformly ^15^N-labeled protein was expressed in *E. coli* BL21 (DE3) cells in M9 minimal medium containing 1.0 g of ^15^NH_4_Cl as the sole nitrogen source. Further expression, isolation and purification of these isotope-labeled proteins was performed as described above for unlabeled protein. To facilitate exchange of deuterated amides back to protons, the purified protein was incubated with 2.5 M guanidinium-HCl and 5 mM EDTA for 3 h, then dialyzed into NMR buffer (10 mM sodium phosphate, pH 7.0, 150 mM NaCl). ^15^N TROSY spectra on samples labeled with only ^15^N were used to confirm nearly complete back-exchange of the deuterated sample. NMR spectra were recorded at 35 °C on a 600 MHz Bruker Avance Neo spectrometer equipped with a cryogenic probe in the METACyt Biomolecular NMR Laboratory at Indiana University, Bloomington.

NMR samples for backbone assignment contained 0.75 mM ^15^N, ^13^C, ^2^H -labeled protein, with or without 0.75 mM ET, in 10 mM sodium phosphate pH 7.0, 150 mM NaCl, and 10% v/v D_2_O, with 0.3 mM 2,2-dimethyl-2-silapentanesulfonic acid (DSS) as an internal reference. Backbone chemical shifts were assigned for each state using TROSY versions of the following standard triple-resonance experiments: HNCACB, HNCOCACB, HNCA, HNCOCA, HNCO, and HNCACO^99^, using non-uniform sampling with Poisson gap schedules ^100^. Data were collected using Topspin 4.1.3 (Bruker). Data were processed using NMRPipe^101^ and istHMS^102^, and analyzed using CARA^103^ and Sparky^104^, all on NMRbox^105^. TALOS-N^106^ was used for chemical shift-based secondary structure predictions^106^. Chemical shift perturbations (CSP) of the backbone upon ligand binding were calculated using ^1^H and ^15^N chemical shifts with Δδ=((ΔδH)^2^+ 0.2(ΔδN)^2^)^1/2^. Chemical shift perturbations upon interaction with L-hercynine were monitored using 0.2 mM ^15^N EgtU and concentrations ranging up to 2 mM. A total of 30 mM betaine was titrated into 0.15 mM ^15^N-labeled EgtU.

### ^15^N spin relaxation experiments

NMR samples for relaxation experiments contained 0.75 mM ^15^N-labeled protein, with or without 0.75 mM ET, in 10 mM sodium phosphate pH 7.0, 150 mM NaCl, and 10% v/v D_2_O, with 0.3 mM 2,2-dimethyl-2-silapentanesulfonic acid (DSS) as an internal reference. The ^15^N spin relaxation rates *R*_1_ and *R*_2_, and ^1^H-^15^N heteronuclear NOE (hNOE) values were measured using TROSY pulse sequences ^107^. The relaxation delays used were 0.05, 0.20, 0.50, 0.80, 1.2, 1.6, 2.0, and 2.5 s for *R*_1_ and 0.017, 0.034, 0.051, 0.068, 0.085, 0.102, 0.119, 0.136, 0.170, and 0.204 s for *R*_2_. Residue-specific *R*_1_ and *R*_2_ values were obtained from fits of peak intensities vs. relaxation time to a single exponential decay function, while hNOE ratios were ascertained directly from intensities in experiments recorded with (2 s relaxation delay followed by 3 s saturation) and without saturation (relaxation delay of 5 s). Errors in hNOE values were calculated by propagating the error from the signal to noise. Hydrogen atoms were added to the crystal structure coordinates for ET-bound EgtU and to the AlphaFold2 model of the apo state using the PDB utilities at http://spin.niddk.nih.gov/bax/nmrserver/pdbutil in order to obtain structure-based predictions for relaxation rates using HYDRONMR^108^. A value of the atomic radius element of 3.8 Å, the known viscosity for water at 35°C ^109^, and CSA of -120 ppm were used for this calculation.

### Ligand specificity analysis using the *Sp*EgtU-SBD-sfGFP titration assay

To measure the ET binding affinity with *Sp*EgtU SBD-sfGFP, the fluorescence change upon ET titration were acquired on a PC1 spectrofluorometer with excitation at 485 nm (2 mm slit) and total emission recorded through a 510 nm cut-off filter in titration buffer (50 mM HEPES, pH 7.5, 150 mM NaCl, 2 mM EDTA) with 2 mM TCEP. ET was titrated into 3 mL 1 µM protein in the same buffer until saturation of the protein was reached. The titration was done with continuous stirring at 25.0 (±0.1) °C and the resulting data fit to a 1:1 protein:ligand binding model to estimate *K*_a_ using DynaFit^95^. The total emission spectrum from 400 nm to 650 nm was measured before and after the titration. The initial emission intensity at 510 nm was set to 1, emission intensity at 650 nm set to 0.

To analyze the ligand specificity of *Sp*EgtU-SBD-sfGFP, triplicate 1 μM protein samples were mixed with 0, 1.0, 10 and 100 μM of the indicated ligand in 100 µL titration buffer (50 mM HEPES, pH 7.5, 150 mM NaCl, 2 mM EDTA) with 2 mM TCEP added in a 96-well plate at 25 ºC. Ligands include *L*-ergothioneine (ET), *L*-hercynine (HER), *L*-histidine (HIS), glycine betaine (GB), proline betaine (PB), choline (CHO), ectoine (ECO), *L*-carnitine (CAR) and dimethylsufoniopropionate (DMSP). Fluorescence was obtained by excitation at 485 nm and emission at 510 nm. After the fluorescence intensity was determined, ET was added into samples to 1.0 µM with 100 μM of the indicated ligand, with the fluorescence intensity of those samples measured again. The change in fluorescence intensity, ΔF, between ET-added samples (Fs) and ET-free samples (Fo) were normalized to the ratio R defined as (|Fo-Fs|)/Fo.

### EFI-GNN analysis

We generated an SSN using the sequence BLAST option with the SBD of the *Sp*EgtU as the query sequence of the UniProt database using the default UniProt BLAST E-value of 5 using the Enzyme Function Institute – Enzyme Similarity Tool (EFI-EST; https://efi.igb.illinois.edu/efi-est/)^73,74^. All of the resulting sequences belonged to the pfam protein family PF04069. and sequences were retrieved in December 2021 using the UniRef90 option. This option takes sequences that share ≥90% sequence identity over 80% of the sequence length, groups them together and represents them by a sequence known as the cluster ID. The resulting sequence file was subjected to SSN analysis using an alignment score of 120 and a minimum and maximum sequence length of 250 and 650 residues in an effort to eliminate truncation artifacts. The resulting SSN was colored and found to contain 19,991 metanodes and 57,649 unique accession IDs that segregate into 2044 non-singleton clusters and 2458 singletons and displayed as a repnode (representative node) 60 file (sequences with 60% identity over 80% of the sequences represented by a single node), analyzed and annotated using Cytoscape^110^. Multiple sequence alignments from each SSN cluster were trimmed for easier visualization using the tool CIAlign^111^ to remove insertions found in fewer than half of the sequences and to crop any poorly aligned termini of sequences. The trimmed multiple sequence alignments were then visualized using WebLogo 3.^112^

### Statistical analysis methods

The number of biological or independent replicates (*n*) is indicated for each experiment and whenever possible all experimental data points are shown along with the standard error of the mean. No statistical method was used to predetermine the sample size.

## ACKNOWLEDGEMENTS

The authors gratefully acknowledge support from the US National Institutes of Health (R35 GM118157). We thank Dr. Brenna Walsh (Indiana University) for synthesis of heavy and light HPE-IAM, Dr. C. M. Pis Diez (Fundación Institut Leloir, Argentina) for early assistance with thiol profiling with HPE-IAM probes, Prof. Malcolm Winkler (Indiana University) for assistance in *S. pneumoniae* D39 strain construction and growth curves, Dr. Hongwei Wu (Indiana University) for help in some of the NMR data acquisition, and Prof. Michelle Reniere (University of Washington) for the gift of *L. monocytogenes* genomic DNA and comments on the manuscript.

## AUTHOR CONTRIBUTIONS

Conceptualization, Y.Z., D.P.G.; Investigation: Y.Z., G.G.-G., K.A.L., K.A.E.; Writing: Y.Z., K.A.E., D.P.G., Funding acquisition: D.P. G., Supervision: Y.Z., K.A.E, D.P.G.

## DATA AVAILABILITY STATEMENT

All primary data are available upon request. The crystallographic structures have been deposited in the Protein Data Bank under accession codes 7TXL and 7TXK. NMR data are available from the BMRB under accession codes 51423 and 51424 for the apo and ET-bound states of the EgtU SBD, respectively. The model of the apo EgtU SBD is available in ModelArchive at https://modelarchive.org/doi/10.5452/ma-xwg27.

## REFERENCES

1. Fahey, R.C. Glutathione analogs in prokaryotes. Biochim Biophys Acta 1830, 3182–98 (2013).

2. Gaballa, A. et al. Biosynthesis and functions of bacillithiol, a major low-molecular-weight thiol in Bacilli. Proc Natl Acad Sci U S A 107, 6482–6486 (2010).

3. Reyes, A.M. et al. Chemistry and Redox Biology of Mycothiol. Antioxid Redox Signal 28, 487–504 (2018).

4. Newton, G.L. & Javor, B. Gamma-Glutamylcysteine and thiosulfate are the major low-molecular-weight thiols in halobacteria. J Bacteriol 161, 438–41 (1985).

5. Ulrich, K. & Jakob, U. The role of thiols in antioxidant systems. Free Radic Biol Med 140, 14–27 (2019).

6. Ung, K.S. & Av-Gay, Y. Mycothiol-dependent mycobacterial response to oxidative stress. FEBS Lett 580, 2712–6 (2006).

7. Reniere, M.L. Reduce, Induce, Thrive: Bacterial Redox Sensing during Pathogenesis. J Bacteriol 200, e00128–18 (2018).

8. Reniere, M.L. et al. Glutathione activates virulence gene expression of an intracellular pathogen. Nature 517, 170–3 (2015).

9. Barger, G. & Ewins, A.J. The constitution of ergothioneine a betaine related to histidine. J Am Chem Soc 99, 2336–2341 (1911).

10. Carlsson, J., Kierstan, M.P. & Brocklehurst, K. Reactions of L-ergothioneine and some other aminothiones with 2, 2′-and 4, 4′-dipyridyl disulphides and of L-ergothioneine with iodoacetamide. 2-Mercaptoimidazoles, 2-and 4-thiopyridones, thiourea and thioacetamide as highly reactive neutral sulphur nucleophiles. Biochem J 139, 221–235 (1974).

11. Fahey, R.C. Glutathione analogs in prokaryotes. Biochim Biophys Acta 1830, 3182–3198 (2013).

12. Jocelyn, P.C. Biochemistry of the SH Group, Academic Press (London) (1972).

13. Long, L.H. & Halliwell, B. Oxidation and generation of hydrogen peroxide by thiol compounds in commonly used cell culture media. Biochem Biophys Res Commun 286, 991–994 (2001).

14. Mayuni, T. et al. Studies on ergothioneine. V. Determination by high performance liquid chromatography and application to metabolic research. Chem Pharmaceut Bull 26, 3772–3778 (1978).

15. Melville, D.B. Ergothioneine. Vitamins & Hormones 17, 155–204 (1959).

16. Misra, H.P. Generation of superoxide free radical during the autoxidation of thiols. J Biol Chem 249, 2151–2155 (1974).

17. Yadan, J.C. Matching chemical properties to molecular biological activities opens a new perspective on L-ergothioneine. FEBS Lett, in the press (doi: 10.1002/1873-3468.14264) (2021).

18. Akanmu, D., Cecchini, R., Aruoma, O.I. & Halliwell, B. The antioxidant action of ergothioneine. Arch Biochem Biophys 288, 10–16 (1991).

19. Aruoma, O.I., Whiteman, M., England, T.G. & Halliwell, B. Antioxidant action of ergothioneine: assessment of its ability to scavenge peroxynitrite. Biochem Biophys Res Commun 231, 389–391 (1997).

20. Hartman, P.E. Ergothioneine as antioxidant. Meth Enzymol 186, 310–318 (1990).

21. Ma, Z. et al. Bacillithiol is a major buffer of the labile zinc pool in Bacillus subtilis. Mol Microbiol 94, 756–770 (2014).

22. Hanlon, D.P. Interaction of ergothioneine with metal ions and metalloenzymes. J Med Chem 14, 1084–1087 (1971).

23. Motohashi, N., Mori, I., Sugiura, Y. & Tanaka, H. Metal complexes of ergothioneine. Chem Pharmaceut Bull 22, 654–657 (1974).

24. Genghof, D.S. Biosynthesis of ergothioneine and hercynine by fungi and Actinomycetales. J Bacteriol 103, 475–478 (1970).

25. Pfeiffer, C., Bauer, T., Surek, B., Schömig, E. & Gründemann, D. Cyanobacteria produce high levels of ergothioneine. Food Chem 129, 1766–1769 (2011).

26. Alamgir, K.M., Masuda, S., Fujitani, Y., Fukuda, F. & Tani, A. Production of ergothioneine by Methylobacterium species. Front Microbiol 6, 1185 (2015).

27. Gamage, A.M. et al. The proteobacterial species Burkholderia pseudomallei produces ergothioneine, which enhances virulence in mammalian infection. FASEB J, fj201800716 (2018).

28. Genghof, D., Inamine, E., Kovalenko, V. & Melville, D. Ergothioneine in microorganisms. J Biol Chem 223, 9–17 (1956).

29. Genghof, D.S. & Damme, O.V. Biosynthesis of ergothioneine and hercynine by mycobacteria. J Bacteriol 87, 852–862 (1964).

30. Melville, D.B. & Eich, S. The occurrence of ergothioneine in plant material. J Biol Chem 18, 647–651 (1956).

31. Stampfli, A.R., Blankenfeldt, W. & Seebeck, F.P. Structural basis of ergothioneine biosynthesis. Curr Opin Struct Biol 65, 1–8 (2020).

32. Stampfli, A.R. & Seebeck, F.P. The catalytic mechanism of sulfoxide synthases. Curr Opin Chem Biol 59, 111–118 (2020).

33. Liao, C. & Seebeck, F.P. Convergent Evolution of Ergothioneine Biosynthesis in Cyanobacteria. Chembiochem 18, 2115–2118 (2017).

34. Seebeck, F.P. In vitro reconstitution of mycobacterial ergothioneine biosynthesis. J Am Chem Soc 132, 6632–3 (2010).

35. Beliaeva, M.A., Leisinger, F. & Seebeck, F.P. In Vitro Reconstitution of a Five-Step Pathway for Bacterial Ergothioneine Catabolism. ACS Chem Biol 16, 397–403 (2021).

36. Cumming, B.M., Chinta, K.C., Reddy, V.P. & Steyn, A.J.C. Role of Ergothioneine in Microbial Physiology and Pathogenesis. Antioxid Redox Signal 28, 431–444 (2018).

37. Saini, V. et al. Ergothioneine maintains redox and bioenergetic homeostasis essential for drug susceptibility and virulence of Mycobacterium tuberculosis. Cell Rep 14, 572–585 (2016).

38. Singh, A.R., Strankman, A., Orkusyan, R., Purwantini, E. & Rawat, M. Lack of mycothiol and ergothioneine induces different protective mechanisms in Mycobacterium smegmatis. Biochem Biophysics Rep 8, 100–106 (2016).

39. Singh, A. et al. Mycobacterium tuberculosis WhiB3 maintains redox homeostasis by regulating virulence lipid anabolism to modulate macrophage response. PLoS Pathog 5, e1000545 (2009).

40. Singh, A. et al. Mycobacterium tuberculosis WhiB3 responds to O2 and nitric oxide via its [4Fe-4S] cluster and is essential for nutrient starvation survival. Proc Natl Acad Sci U S A 104, 11562–11567 (2007).

41. Richard-Greenblatt, M. et al. Regulation of ergothioneine biosynthesis and its effect on Mycobacterium tuberculosis growth and infectivity. J Biol Chem 290, 23064–23076 (2015).

42. Bello, M.H., Barrera-Perez, V., Morin, D. & Epstein, L. The Neurospora crassa mutant NcΔEgt-1 identifies an ergothioneine biosynthetic gene and demonstrates that ergothioneine enhances conidial survival and protects against peroxide toxicity during conidial germination. Fung Gen Biol 49, 160–172 (2012).

43. Bello, M.H., Mogannam, J.C., Morin, D. & Epstein, L. Endogenous ergothioneine is required for wild type levels of conidiogenesis and conidial survival but does not protect against 254 nm UV-induced mutagenesis or kill. Fung Gen Biol 73, 120–127 (2014).

44. Gallagher, L. et al. The Aspergillus fumigatus protein GliK protects against oxidative stress and is essential for gliotoxin biosynthesis. Eukaryot Cell 11, 1226–38 (2012).

45. Sheridan, K.J. et al. Ergothioneine Biosynthesis and Functionality in the Opportunistic Fungal Pathogen, Aspergillus fumigatus. Sci Rep 6, 35306 (2016).

46. Pluskal, T., Ueno, M. & Yanagida, M. Genetic and Metabolomic Dissection of the Ergothioneine and Selenoneine Biosynthetic Pathway in the Fission Yeast, S. pombe, and Construction of an Overproduction System. PLoS One 9, e97774 (2014).

47. Cheah, I.K. & Halliwell, B. Ergothioneine, recent developments. Redox Biol 42, 101868 (2021).

48. Grundemann, D., Hartmann, L. & Flogel, S. The ergothioneine transporter (ETT): Substrates and locations, an inventory. FEBS Lett, in the press (doi: 10.1002/1873-3468.14269) (2021).

49. Grundemann, D. et al. Discovery of the ergothioneine transporter. Proc Natl Acad Sci U S A 102, 5256–61 (2005).

50. Tamai, I. et al. Cloning and characterization of a novel human pH-dependent organic cation transporter, OCTN1. FEBS Lett 419, 107–11 (1997).

51. Nikodemus, D. et al. Paramount levels of ergothioneine transporter SLC22A4 mRNA in boar seminal vesicles and cross-species analysis of ergothioneine and glutathione in seminal plasma. J Physiol Pharmacol 62, 411–9 (2011).

52. Tokuhiro, S. et al. An intronic SNP in a RUNX1 binding site of SLC22A4, encoding an organic cation transporter, is associated with rheumatoid arthritis. Nat Genet 35, 341–8 (2003).

53. Harwood, M.D., Zhang, M., Pathak, S.M. & Neuhoff, S. The Regional-Specific Relative and Absolute Expression of Gut Transporters in Adult Caucasians: A Meta-Analysis. Drug Metab Dispos 47, 854–864 (2019).

54. Monaco, G. et al. RNA-Seq Signatures Normalized by mRNA Abundance Allow Absolute Deconvolution of Human Immune Cell Types. Cell Rep 26, 1627–1640 e7 (2019).

55. Berg, T. et al. Expression of MATE1, P-gp, OCTN1 and OCTN2, in epithelial and immune cells in the lung of COPD and healthy individuals. Respir Res 19, 68 (2018).

56. Sleator, R.D., Wemekamp-Kamphuis, H.H., Gahan, C.G., Abee, T. & Hill, C. A PrfA-regulated bile exclusion system (BilE) is a novel virulence factor in Listeria monocytogenes. Mol Microbiol 55, 1183–95 (2005).

57. Zhang, Y., Martin, J.E., Edmonds, K.A., Winkler, M.E. & Giedroc, D.P. SifR is an Rrf2-family quinone sensor associated with catechol iron uptake in Streptococcus pneumoniae D39. J Biol Chem, accepted for publication (2022).

58. Capdevila, D.A. et al. Tuning site-specific dynamics to drive allosteric activation in a pneumococcal zinc uptake regulator. Elife 7, e37268 (2018).

59. Beinker, P. et al. Crystal structures of SnoaL2 and AclR: Two putative hydroxylases in the biosynthesis of aromatic polyketide antibiotics. J Mol Biol 359, 728–40 (2006).

60. Horn, C. et al. Molecular determinants for substrate specificity of the ligand-binding protein OpuAC from Bacillus subtilis for the compatible solutes glycine betaine and proline betaine. J Mol Biol 357, 592–606 (2006).

61. Potter, A.J., Trappetti, C. & Paton, J.C. Streptococcus pneumoniae uses glutathione to defend against oxidative stress and metal ion toxicity. J Bacteriol 194, 6248–54 (2012).

62. Ruiz, S.J., Schuurman-Wolters, G.K. & Poolman, B. Crystal structure of the substrate-binding domain from Listeria monocytogenes bile-resistance determinant BilE. Crystals 6, 162 (2016).

63. Quiocho, F.A. & Ledvina, P.S. Atomic structure and specificity of bacterial periplasmic receptors for active transport and chemotaxis: Variation of common themes. Mol Microbiol 20, 17–25 (1996).

64. Chu, B.C. & Vogel, H.J. A structural and functional analysis of type III periplasmic and substrate binding proteins: their role in bacterial siderophore and heme transport. Biol Chem 392, 39–52 (2011).

65. Berntsson, R.P., Smits, S.H., Schmitt, L., Slotboom, D.J. & Poolman, B. A structural classification of substrate-binding proteins. FEBS Lett 584, 2606–17 (2010).

66. Schiefner, A., Holtmann, G., Diederichs, K., Welte, W. & Bremer, E. Structural basis for the binding of compatible solutes by ProX from the hyperthermophilic archaeon Archaeoglobus fulgidus. J Biol Chem 279, 48270–81 (2004).

67. Pittelkow, M., Tschapek, B., Smits, S.H., Schmitt, L. & Bremer, E. The crystal structure of the substrate-binding protein OpuBC from Bacillus subtilis in complex with choline. J Mol Biol 411, 53–67 (2011).

68. Tame, J.R., Sleigh, S.H., Wilkinson, A.J. & Ladbury, J.E. The role of water in sequence-independent ligand binding by an oligopeptide transporter protein. Nat Struct Biol 3, 998–1001 (1996).

69. Baker, B.M. & Murphy, K.P. Prediction of binding energetics from structure using empirical parameterization. Methods Enzymol 295, 294–315 (1998).

70. Qu, F., ElOmari, K., Wagner, A., De Simone, A. & Beis, K. Desolvation of the substrate-binding protein TauA dictates ligand specificity for the alkanesulfonate ABC importer TauABC. Biochem J 476, 3649–3660 (2019).

71. Jumper, J. et al. Highly accurate protein structure prediction with AlphaFold. Nature 596, 583–589 (2021).

72. Smirnova, I., Kasho, V. & Kaback, H.R. Real-time conformational changes in LacY. Proc Natl Acad Sci U S A 111, 8440–5 (2014).

73. Gerlt, J.A. Genomic Enzymology: Web Tools for Leveraging Protein Family Sequence-Function Space and Genome Context to Discover Novel Functions. Biochemistry 56, 4293–4308 (2017).

74. Zallot, R., Oberg, N. & Gerlt, J.A. The EFI Web Resource for Genomic Enzymology Tools: Leveraging Protein, Genome, and Metagenome Databases to Discover Novel Enzymes and Metabolic Pathways. Biochemistry 58, 4169–4182 (2019).

75. Teichmann, L., Kummel, H., Warmbold, B. & Bremer, E. OpuF, A New Bacillus Compatible Solute ABC Transporter with a Substrate-Binding Protein Fused to the Transmembrane Domain. Appl Environ Microbiol 84, e01728–18 (2018).

76. Michel, A.M. et al. Cellular adaptation of Clostridioides difficile to high salinity encompasses a compatible solute-responsive change in cell morphology. Environ Microbiol 24, 1499–1517 (2022).

77. Cheah, I.K., Lee, J.Z., Tang, R.M.Y., Koh, P.W. & Halliwell, B. Does Lactobacillus reuteri influence ergothioneine levels in the human body? FEBS Lett, in the press (doi: 10.1002/1873-3468.14364) (2022).

78. Schnizlein, M.K. & Young, V.B. Capturing the environment of the Clostridioides difficile infection cycle. Nat Rev Gastroenterol Hepatol, in the press (doi: 0.1038/s41575-022-00610-0) (2022).

79. Saini, V. et al. Ergothioneine Maintains Redox and Bioenergetic Homeostasis Essential for Drug Susceptibility and Virulence of Mycobacterium tuberculosis. Cell Rep 14, 572–85 (2016).

80. Slager, J., Aprianto, R. & Veening, J.W. Deep genome annotation of the opportunistic human pathogen Streptococcus pneumoniae D39. Nucleic Acids Res 46, 9971–9989 (2018).

81. Palazzolo-Ballance, A.M. et al. Neutrophil microbicides induce a pathogen survival response in community-associated methicillin-resistant Staphylococcus aureus. J Immunol 180, 500–509 (2008).

82. Lengfelder, I. et al. Complex Bacterial Consortia Reprogram the Colitogenic Activity of Enterococcus faecalis in a Gnotobiotic Mouse Model of Chronic, Immune-Mediated Colitis. Front Immunol 10, 1420 (2019).

83. Cheah, I.K., Tang, R.M., Yew, T.S., Lim, K.H. & Halliwell, B. Administration of Pure Ergothioneine to Healthy Human Subjects: Uptake, Metabolism, and Effects on Biomarkers of Oxidative Damage and Inflammation. Antioxid Redox Signal 26, 193–206 (2017).

84. Goncharenko, K.V. et al. Selenocysteine as a Substrate, an Inhibitor and a Mechanistic Probe for Bacterial and Fungal Iron-Dependent Sulfoxide Synthases. Chemistry 26, 1328–1334 (2019).

85. Lanie, J.A. et al. Genome sequence of Avery’s virulent serotype 2 strain D39 of Streptococcus pneumoniae and comparison with that of unencapsulated laboratory strain R6. J Bacteriol 189, 38–51 (2007).

86. Zhang, Y. et al. The Pneumococcal Iron Uptake Protein A (PiuA) Specifically Recognizes Tetradentate Fe(III) bis-and Mono-Catechol Complexes. J Mol Biol 432, 5390–5410 (2020).

87. Ducret, A., Quardokus, E.M. & Brun, Y.V. MicrobeJ, a tool for high throughput bacterial cell detection and quantitative analysis. Nat Microbiol 1, 16077 (2016).

88. Nogales, E., Downing, K.H., Amos, L.A. & Lowe, J. Tubulin and FtsZ form a distinct family of GTPases. Nat Struct Biol 5, 451–8 (1998).

89. Hamid, H.A. et al. Polysulfide stabilization by tyrosine and hydroxyphenyl-containing derivatives that is important for a reactive sulfur metabolomics analysis. Redox Biol 21, 101096 (2019).

90. Walsh, B.J.C. et al. The Response of Acinetobacter baumannii to Hydrogen Sulfide Reveals Two Independent Persulfide-Sensing Systems and a Connection to Biofilm Regulation. mBio 11, e01254–20 (2020).

91. Peroutka III, R.J., Orcutt, S.J., Strickler, J.E. & Butt, T.R. SUMO fusion technology for enhanced protein expression and purification in prokaryotes and eukaryotes. Methods Mol Biol 705, 15–30 (2011).

92. Nichols, A.L. et al. Fluorescence activation mechanism and imaging of drug permeation with new sensors for smoking-cessation ligands. Elife 11, e74648 (2022).

93. Perez, A.J. et al. Movement dynamics of divisome proteins and PBP2x:FtsW in cells of Streptococcus pneumoniae. Proc Natl Acad Sci U S A 116, 3211–3220 (2019).

94. Gibson, D.G. et al. Enzymatic assembly of DNA molecules up to several hundred kilobases. Nat Methods 6, 343–5 (2009).

95. Kuzmic, P. Program DYNAFIT for the analysis of enzyme kinetic data: application to HIV proteinase. Anal Biochem 237, 260–273 (1996).

96. Kabsch, W. Automatic Processing of Rotation Diffraction Data from Crystals of Initially Unknown Symmetry and Cell Constants. J Appl Crystallogr 26, 795–800 (1993).

97. Echols, N. et al. Graphical tools for macromolecular crystallography in PHENIX. J Appl Crystallogr 45, 581–586 (2012).

98. Headd, J.J. et al. Use of knowledge-based restraints in phenix.refine to improve macromolecular refinement at low resolution. Acta Crystallogr 68, 381–90 (2012).

99. Salzmann, M., Wider, G., Pervushin, K. & Wuthrich, K. Improved sensitivity and coherence selection for [15N,1H]-TROSY elements in triple resonance experiments. J Biomol NMR 15, 181–184 (1999).

100. Hyberts, S.G., Takeuchi, K. & Wagner, G. Poisson-gap sampling and forward maximum entropy reconstruction for enhancing the resolution and sensitivity of protein NMR data. J Am Chem Soc 132, 2145–7 (2010).

101. Delaglio, F. et al. NMRPipe: a multidimensional spectral processing system based on UNIX pipes. J Biomol NMR 6, 277–293 (1995).

102. Hyberts, S.G., Milbradt, A.G., Wagner, A.B., Arthanari, H. & Wagner, G. Application of iterative soft thresholding for fast reconstruction of NMR data non-uniformly sampled with multidimensional Poisson Gap scheduling. J Biomol NMR 52, 315–27 (2012).

103. Keller, R.L.J. The Computer Aided Resonance Assignment Tutorial. http://www.nmr.ch (Cantina Verlag, Goldau, 2004).

104. Lee, W., Tonelli, M. & Markley, J.L. NMRFAM-SPARKY: enhanced software for biomolecular NMR spectroscopy. Bioinformatics 31, 1325–7 (2015).

105. Maciejewski, M.W. et al. NMRbox: A Resource for Biomolecular NMR Computation. Biophys J 112, 1529–1534 (2017).

106. Shen, Y. & Bax, A. Protein backbone and sidechain torsion angles predicted from NMR chemical shifts using artificial neural networks. J Biomol NMR 56, 227–41 (2013).

107. Zhu, G., Xia, Y., Nicholson, L.K. & Sze, K.H. Protein dynamics measurements by TROSY-based NMR experiments. J Magn Reson 143, 423–6 (2000).

108. Garcia de la Torre, J., Huertas, M.L. & Carrasco, B. HYDRONMR: Prediction of NMR relaxation of globular proteins from atomic-level structures and hydrodynamic calculations. J Magn Reson 147, 138–46 (2000).

109. Cho, C.H., Urquidi, J., Singh, S. & Robinson, G.W. Thermal offset viscosities of liquid H2O, D2O, and T2O. J Phys Chem B 103, 1991–1994 (1999).

110. Shannon, P. et al. Cytoscape: a software environment for integrated models of biomolecular interaction networks. Genome Res 13, 2498–504 (2003).

111. Tumescheit, C., Firth, A.E. & Brown, K. CIAlign - A highly customisable command line tool to clean, interpret and visualise multiple sequence alignments. Biorxiv, doi: https://doi.org/10.1101/2020.09.14.291484 (2021).

112. Crooks, G.E., Hon, G., Chandonia, J.M. & Brenner, S.E. WebLogo: a sequence logo generator. Genome Res 14, 1188–90 (2004).

