## Supporting Information for "Discovery and structure of a widespread bacterial ABC transporter specific for ergothioneine"

This file contains **Supplementary Figures 1-15**, **Supplemental Tables 1-5** and Supplemental References

**Table of Contents**


| Item | Page |
|---|---|
| **Supplementary Figure 1.** Chemical structures and exact masses of HPE-IAM-derivatized thiols used in this work……………………………………………………...... | S3 |
| **Supplementary Figure 2.** Mass spectrometry of light ($H_4$) and heavy ($d_4$) HPE-IAM-capped thiol standards……………………………………………………....................... | S4 |
| **Supplementary Figure 3.** Mass spectrometry of light ($H_4$) and heavy ($d_4$) HPE-IAM-capped thiols obtained from *S. pneumoniae* D39 cell lysates grown on BHI………….. | S5 |
| **Supplementary Figure 4.** Mass spectrometry of light ($H_4$) and heavy ($d_4$) HPE-IAM-capped thiols obtained from *S. pneumoniae* D39 cell lysates grown on a chemically defined growth medium (CDM) to which exogenous ET was added……........................ | S6 |
| **Supplementary Figure 5**. Replacement and truncation of the C-terminal five residues of *Sp*EgtU SBD (GLLKK) with VC has a minimal effect on the structure and no impact on ET-binding affinity………………………………………………………….. | S7 |
| **Supplementary Figure 6**. Conformational changes and dynamics of *Sp*EgtU SBD in solution. ……………………………………………………………............................... | S8 |
| **Supplementary Figure 7.** Backbone $^1$H,$^{15}$N assignments of apo *Sp*EgtU SBD……… | S9 |
| **Supplementary Figure 8.** Backbone $^1$H,$^{15}$N assignments of ET-bound *Sp*EgtU SBD………………………………………………………..………………………………… | S10 |
| **Supplementary Figure 9.** Chemical structures of QAC ligands used in this work and representative ITC titrations of wild-type *Sp*EgtU SBD with non-cognate ligands…….... | S11 |
| **Supplementary Figure 10.** NMR-monitored titrations of SpEgtU SBD with non-cognate and weakly binding ligands *L*-hercynine and glycine-betaine………………….. | S12 |
| **Supplementary Figure 11**. Sequence similarity network (SSN) of solute binding protein (domains) most closely related to *Sp*EgtU SBD………………………………….. | S13 |
| **Supplementary Figure 12.** Network connectivity analysis……………………………… | S14 |
| **Supplementary Figure 13.** The biological range determined for the EgtU subcluster of cluster 2. …………………………………………………….......................................... | S15 |
| **Supplementary Figure 14.** Sequence conservation maps of SSN cluster 2 subclusters…..………………………………………………………...................................... | S16 |
| **Supplementary Figure 15.** ET-binding properties of candidate EgtU homologs from other firmicutes. …………………………………………………….............................. | S17 |
| **Supplementary Table 1.** Data collection and refinement statistics…………………….. | S18 |
| **Supplementary Table 2.** List of water molecules and associated B-factors in the structures of *Sp*EgtU $SBD_{CTT}$ and *Sp*EgtU SBD…………………………………………… | S19 |
| **Supplementary Table 3.** List of *Streptococcus pneumoniae* D39 strains in this work.. | S20 |
| **Supplementary Table 4.** Primers used in this study……………………………………... | S20 |
| **Supplementary Table 5.** Molar extinction coefficients of purified proteins at 280 nm ($\varepsilon_{280}$) used in this work……………………………………………………........................ | S21 |
| Supplementary References………………………………………………………................. | S21 |

**SUPPLEMENTARY FIGURES**

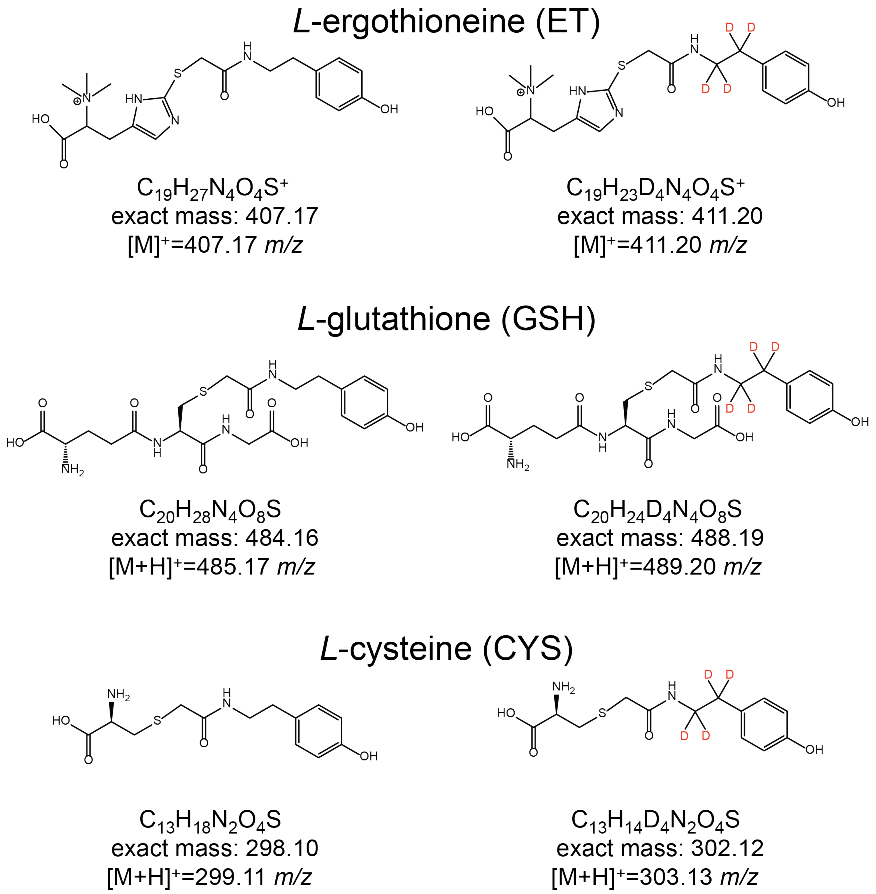


**Supplementary Figure 1. Chemical structures and exact masses of HPE-IAM-derivatized thiols used in this work.** The light ($H_4$; *left*) and heavy ($d_4$; *right*) HPE-IAM adducts of *L*-ergothioneine (*top*), *L*-glutathione (*middle*) and *L*-cysteine (*bottom*).

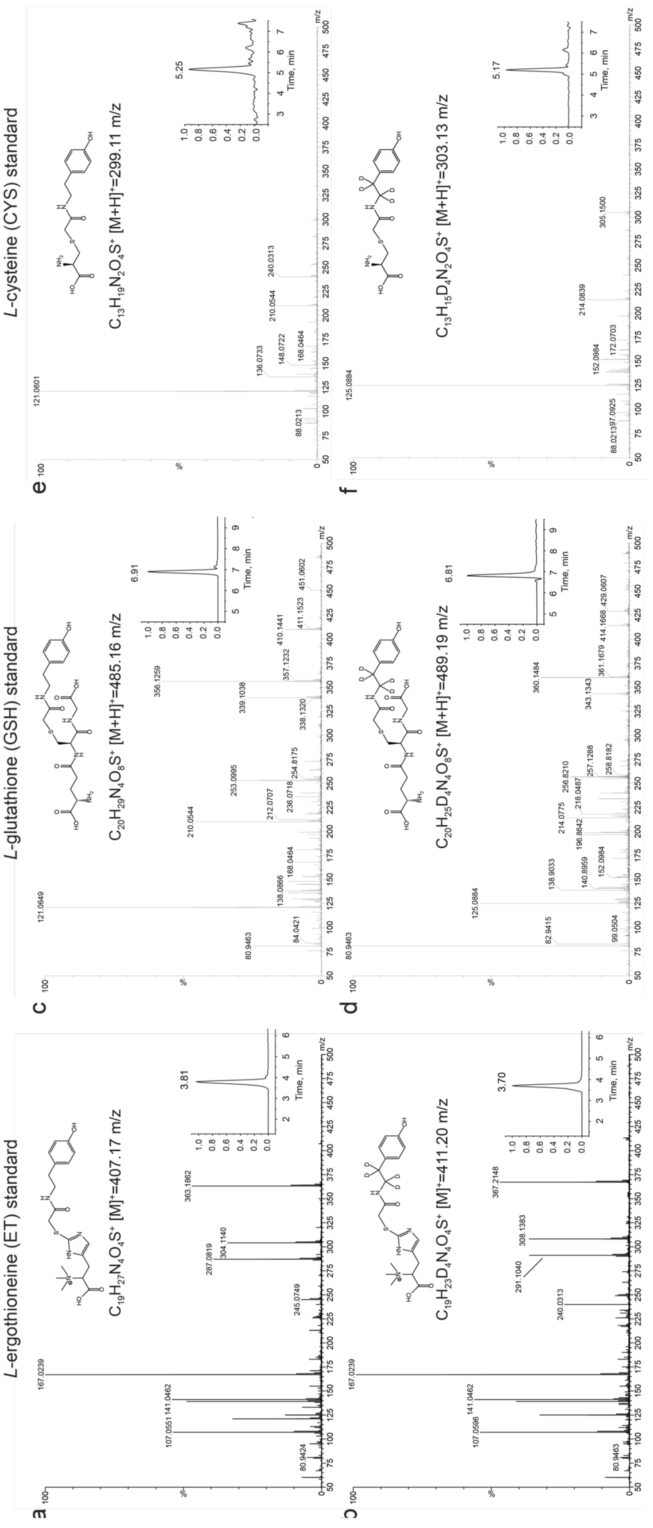


**Supplementary Figure 2. Mass spectrometry of light ($H_4$) and heavy (*$d_4$*) HPE-IAM-capped thiol standards.** LC-MS (MS1) (*insets*) and LC-MS/MS (normalized ion count) of authentic HPE-IAM capped LMW thiols are shown. **a** and **b**, *L*-ergothioneine; **c** and **d**, *L*-glutathione; **e** and **f**, *L*-cysteine. Structure of the compound and expected masses used to query the TIC are shown.

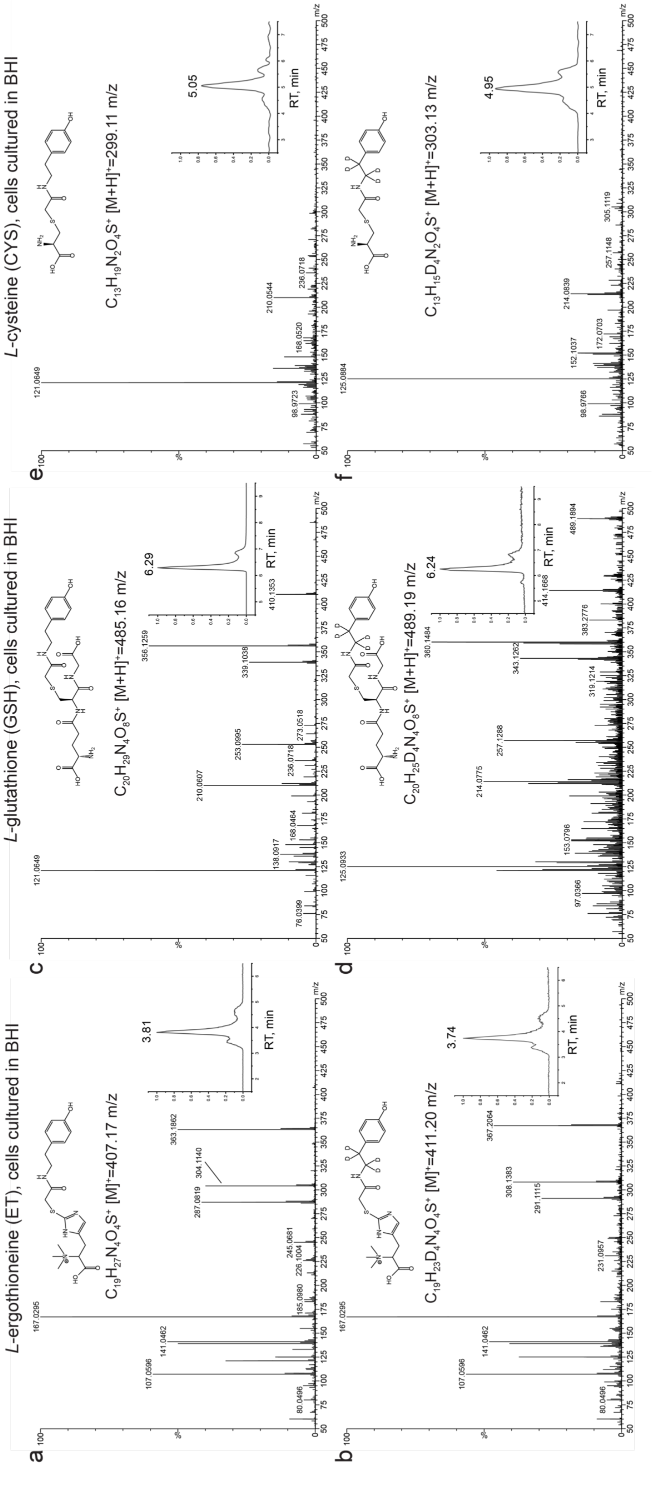


**Supplementary Figure 3. Mass spectrometry of light ($H_4$) and heavy (*$d_4$*) HPE-IAM-capped thiols obtained from *S. pneumoniae* D39 cell lysates grown on BHI.** LC-MS (MS1) (*insets*) and LC-MS/MS (normalized ion count) of HPE-IAM capped LMW thiols are shown. **a** and **b**, *L*-ergothioneine; **c** and **d**, *L*-glutathione; **e** and **f**, *L*-cysteine. Structure of the compound and expected masses used to query the TIC are shown.

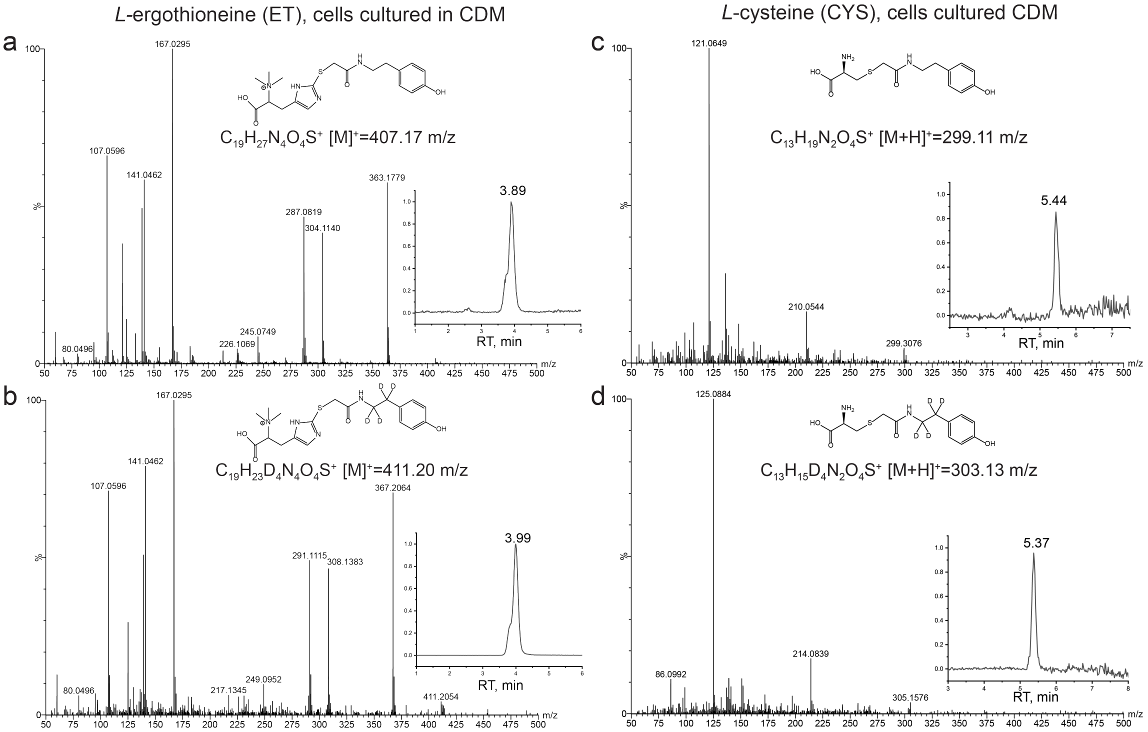


**Supplementary Figure 4. Mass spectrometry of light ($H_4$) and heavy ($d_4$) HPE-IAM-capped thiols obtained from *S. pneumoniae* D39 cell lysates grown on a chemically defined growth medium (CDM) to which exogenous ET was added.** LC-MS (MS1) (*insets*) and LC-MS/MS (normalized ion count) of HPE-IAM capped LMW thiols are shown. **a** and **b**, *L*-ergothioneine; **c** and **d**, *L*-cysteine. Structure of the compound and expected masses used to query the TIC are shown. Note that there is no detectable *L*-glutathione in these lysates.

.

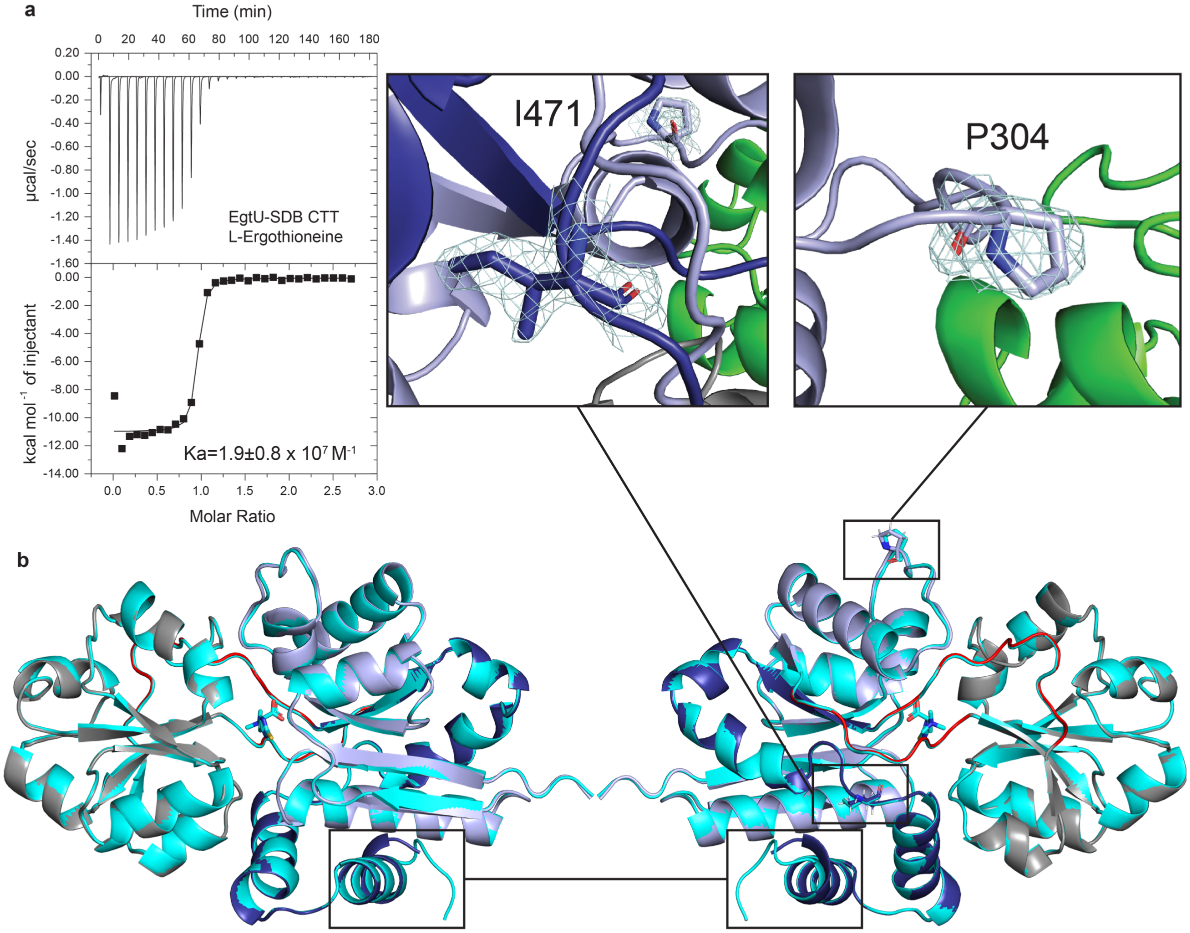


**Supplementary Figure 5. Replacement and truncation of the C-terminal five residues of SpEgtU SBD (GLLKK) with VC has a minimal effect on the structure and no impact on ET-binding affinity. a**, Isothermal titration calorimetry of the C-terminally truncated *Sp*EgtU SBD, designated *Sp*EgtU $SBD_{CTT}$, titrated with ET. The continuous line through the EgtU $SBD_{CTT}$ binding data represents a fit to a 1:1 binding model (see Table 1, main text for parameters). **b**, Global alignment of the crystal structures of ET-bound *Sp*EgtU SBD (*cyan*, PDB code: 7TXK) and *Sp*EgtU $SBD_{CTT}$ (*gray, light blue, and dark blue,* PDB code:7TXL) in cartoon representation (see Supplementary Table 1 for structure statistics). ET is shown as cyan sticks. The structure of the C-terminus is boxed. The expanded regions of the structure shown in the boxes (*top*) highlight an unusual rotamer for I471 side chain and the *cis* conformation of P304 side chain.

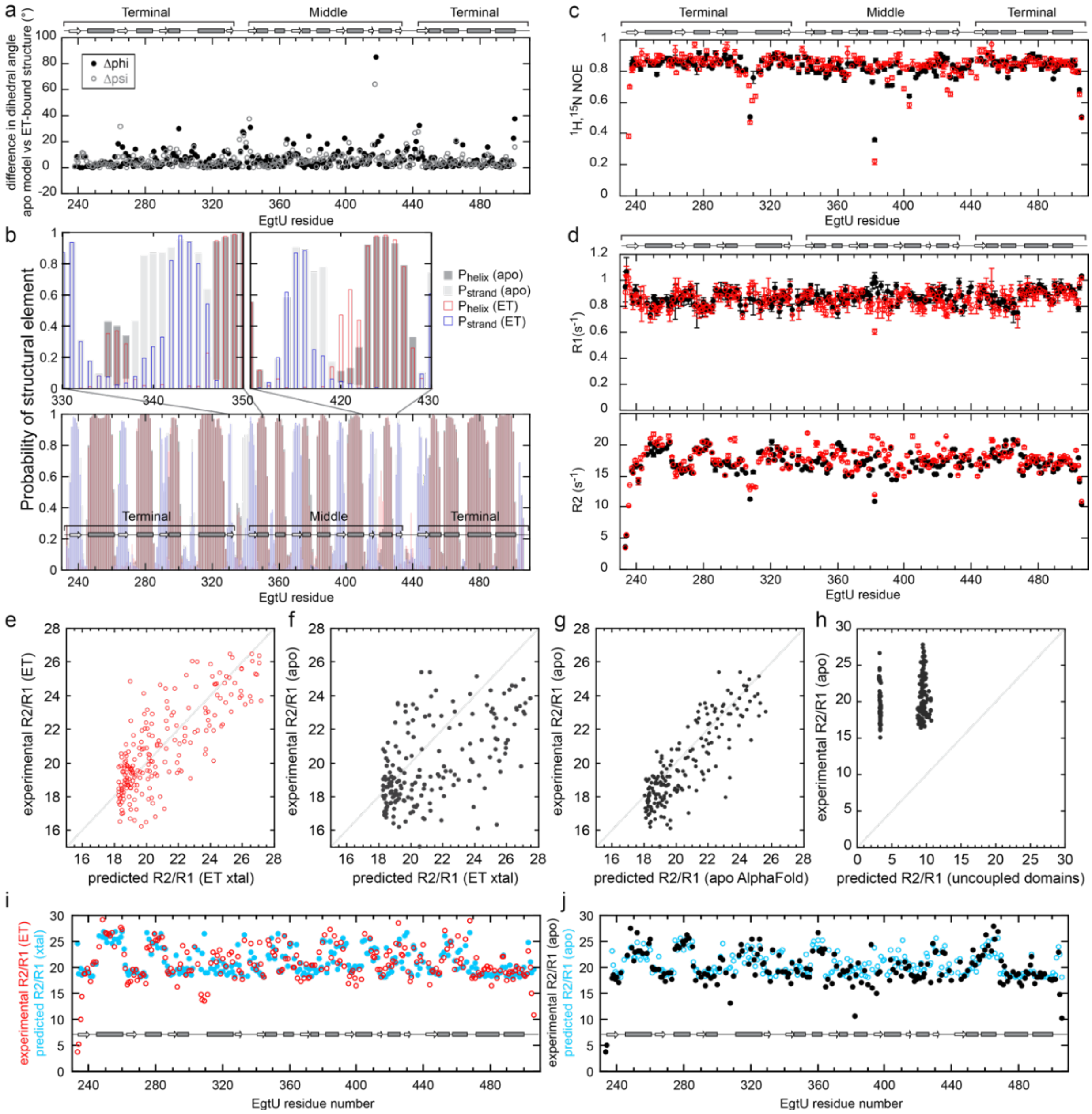


**Supplementary Figure 6. Conformational changes and dynamics of *Sp*EgtU SBD in solution. a,** Differences in backbone dihedral angles phi and psi between the AlphaFold2 model of apo EgtU-SBD and the crystal structure of ET-bound EgtU-SBD, for each residue predict limited conformational changes localized to linkers and loops **b,** Chemical shift-based secondary structure predictions for EgtU in the apo and ET-bound states in solution shows extended β strands near the hinge in the apo state. **c.** Steady-state heteronuclear $^{15}$N[$^{1}$H] NOE reporting on sub-nanosecond amide bond vector motions of for apo EgtU SBD (black circles) and ET-bound EgtU SBD (red circles) **d.** Backbone $^{15}$N longitudinal (top) and transverse (bottom) spin relaxation rates of EgtU SBD in the apo (black circles) and ET-bound (red circles) forms. **e.** Comparison between experimental and HYDRONMR calculated values of R2/R1 for ET-bound EgtU-SBD based on the WT crystal structure. **f-h**. Comparison between experimental and HYDRONMR calculated values of R2/R1 for apo EgtU-SBD based on three different models: **f.** an apo structure identical to the ET-bound crystal structure, **g.** an AlphaFold2 model of the open state, and **h.** a model of independently tumbling middle and terminal domains with an infinitely flexible linker. **i.** Experimental (red) and predicted (blue) R2/R1 values for ET-bound $_{15}$N EgtU SBD based on the WT crystal structure. **j.** Experimental (black) and predicted (blue) R2/R1 values for apo $^{15}$N EgtU SBD based on the AlphaFold2 model.

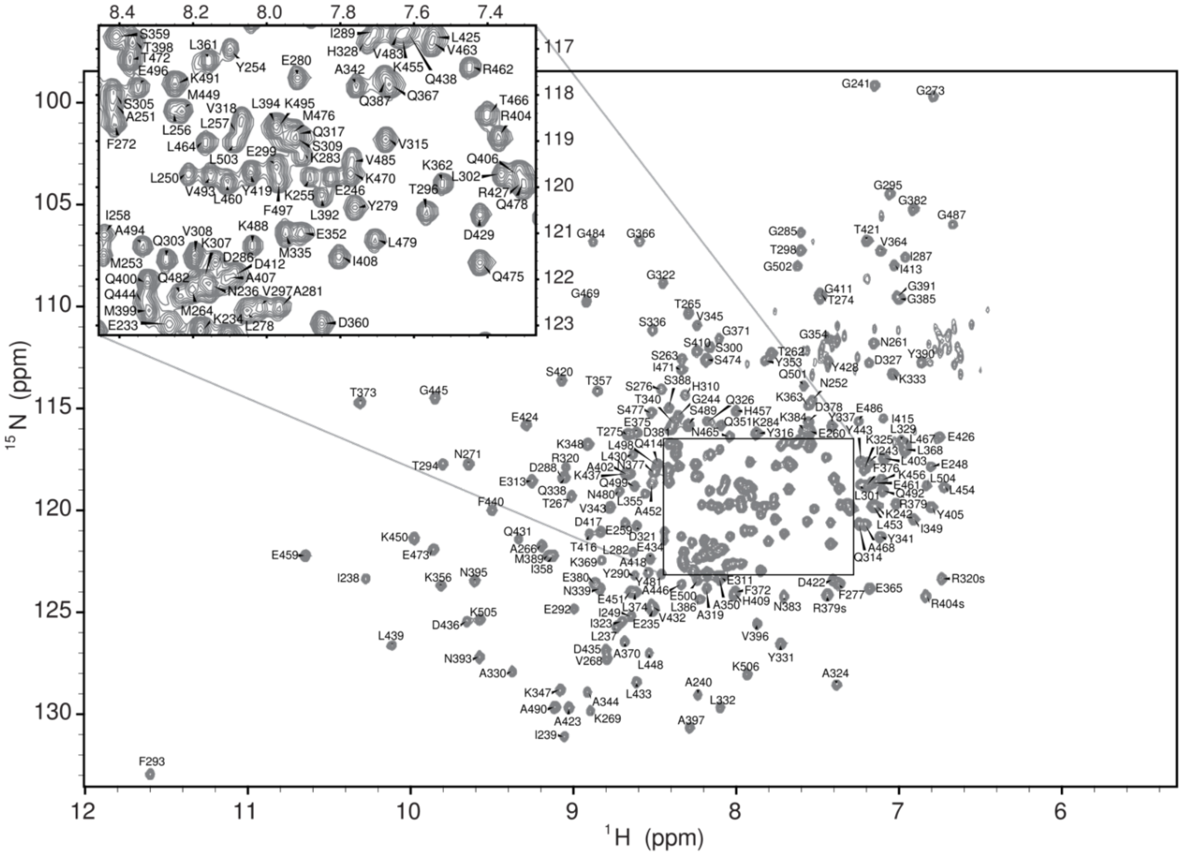


**Supplementary Figure 7**. Backbone $^{1}$H,$^{15}$N assignments of apo *Sp*EgtU SBD shown on an $^{1}$H,$^{15}$N TROSY spectrum of $^{2}$H,$^{13}$C,$^{15}$N-labeled *Sp*EgtU SBD.

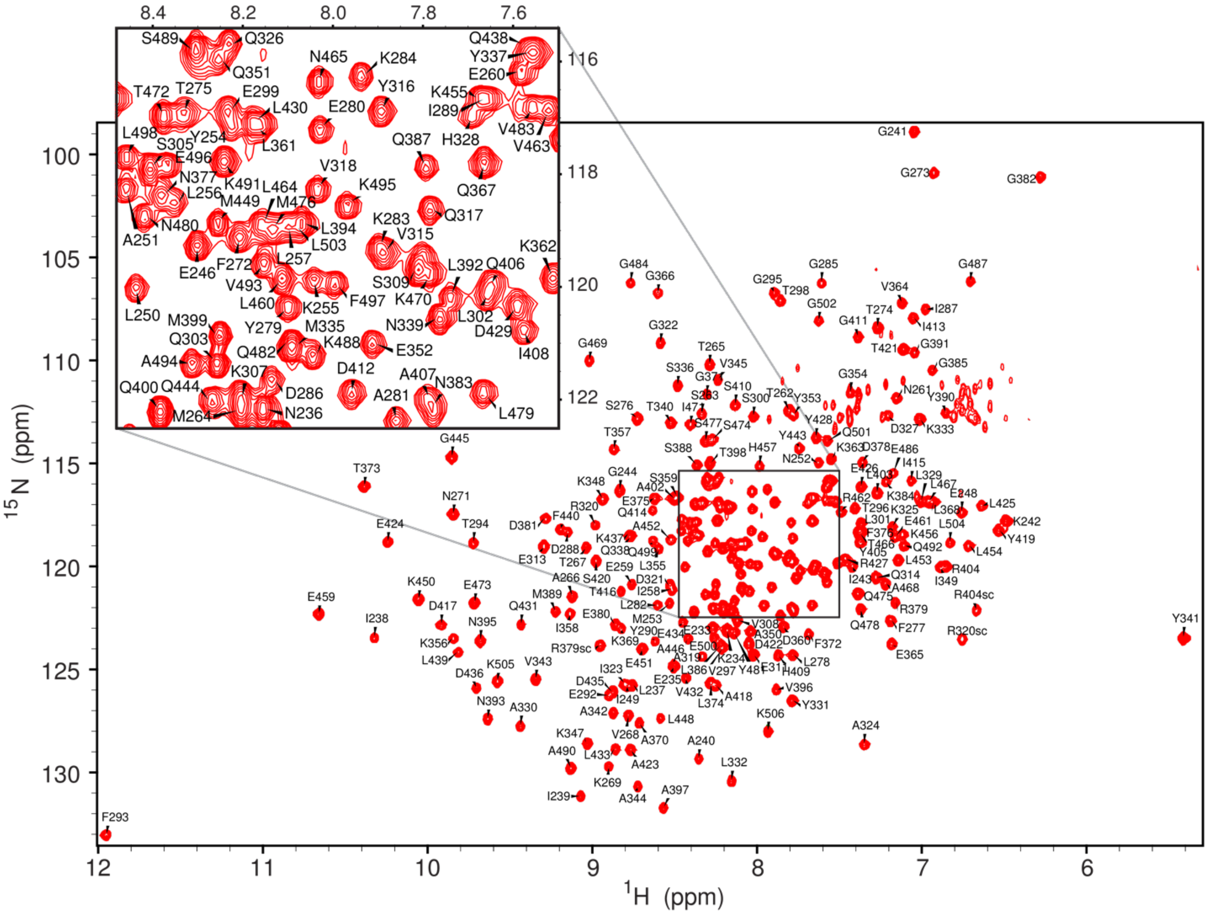


**Supplementary Figure** . Backbone $^{1}$H,$^{15}$N assignments of ET-bound *Sp*EgtU SBD shown on an $^{1}$H,$^{15}$N TROSY spectrum of $^{2}$H,$^{13}$C,$^{15}$N-labeled *Sp*EgtU SBD.

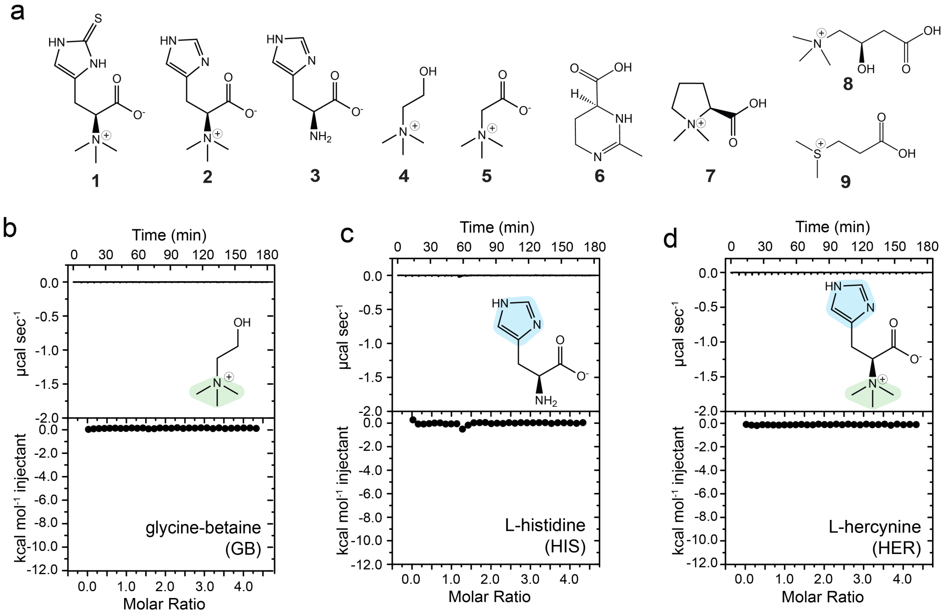


**Supplementary Figure 9. Chemical structures of QAC ligands used in this work and representative ITC titrations of wild-type *Sp*EgtU SBD with non-cognate ligands. a**, QAC Ligands used in this work. **1,** *L*-ergothioneine (ET); **2,** *L*-hercynine (HER); **3,** *L*-histidine (HIS); **4,** choline (CHO); **5,** glycine-betaine (GB); **6,** ectoine (ECT); **7,** proline-betaine (PB); **8,** *L*-carnitine (CAR); **9,** dimethypropiothetin hydrochloride (DMSP). **b-d**, Isothermal titration calorimetry of the wild-type *Sp*EgtU SBD titrated with glycine-betaine (**b**), *L*-histidine (**c)** and *L*-hercynine (**d**) under the same conditions as described in the main text for ET titrations (Fig. 3).

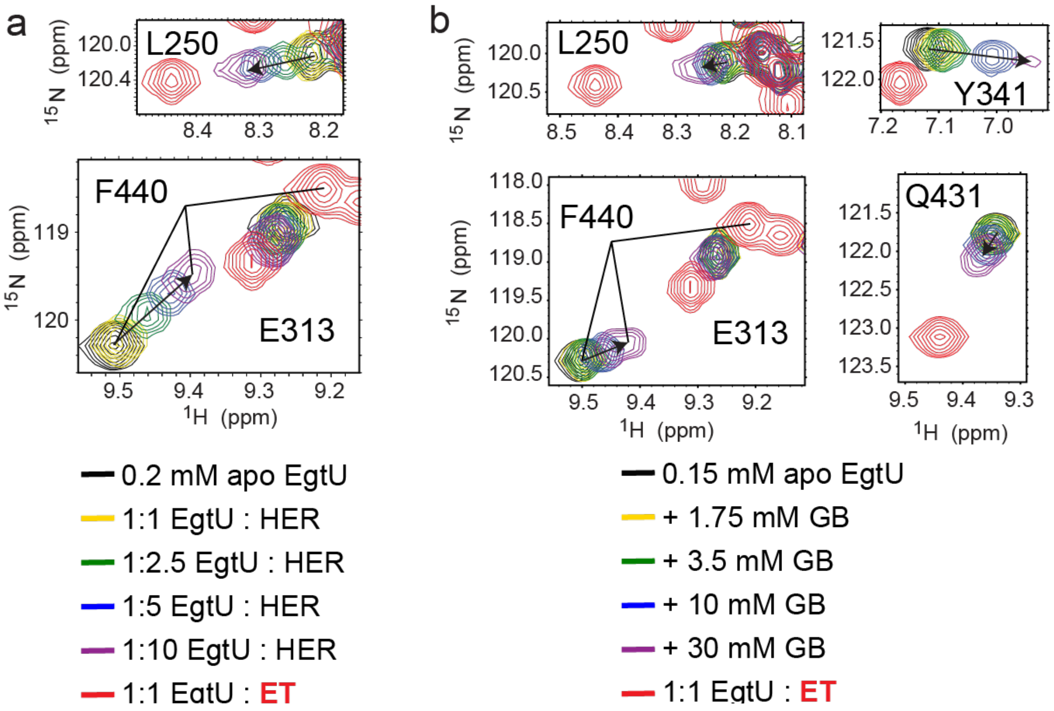


**Supplementary Figure 10. NMR-monitored titrations of *Sp*EgtU SBD with non-cognate and weakly binding ligands *L*-hercynine and glycine betaine. a**, Movement of the indicated backbone NH crosspeak from the apo-state (black) as *L*-hercynine (HER) is added (yellow to purple), compared to the crosspeak position of ET-bound EgtU SBD (red) shown for reference. **b**, Movement of the indicated backbone NH crosspeak from the apo-state (black) as glycine-betaine (GB) is added (yellow to purple), compared to the crosspeak position of ET-bound EgtU SBD (red) shown for reference.

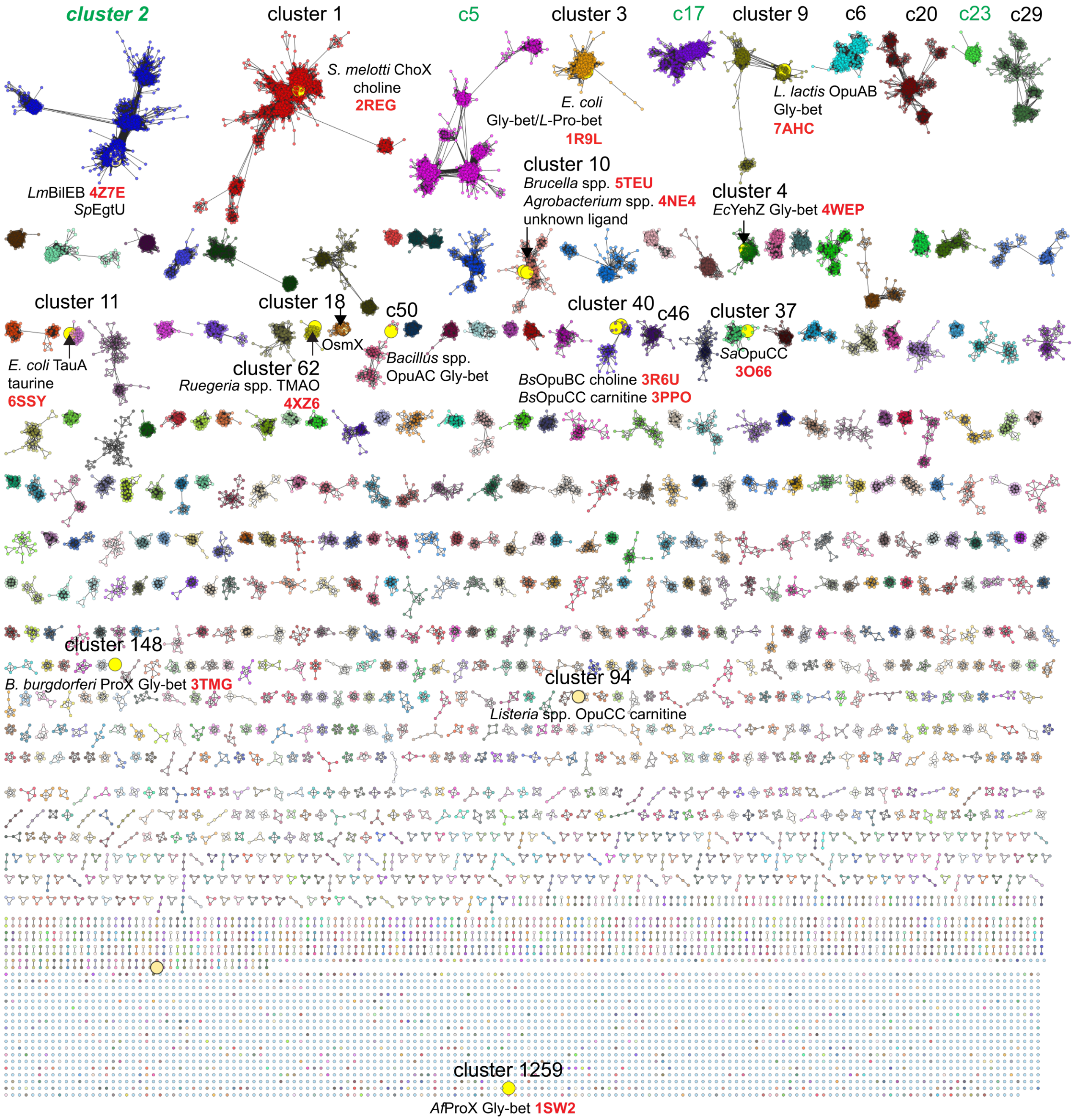


**Supplementary Figure 11**. **Sequence similarity network (SSN) of solute binding protein (domains) most closely related to *Sp*EgtU SBD.** Full sequence similarity network (SSN) analysis[1] of the osmoprotectant subclass of solute binding proteins (or domains) (SBPs/SPDs) resulting in 2,044 clusters and the 2458 singletons. Clusters are ranked numerically from that containing the largest (cluster 1; c1) to the smallest number of sequences and are arranged from *upper left* to *lower right* on the basis of metanode cluster count. Those sequences for which SwissProt annotations exist are highlighted (*yellow circle*), along with their trivial names, known or inferred ligand specificity and the organism (*Sp*, *S. pneumoniae*; *Lm*, *Listeria monocytogenes*, *Bs*, *B. subtilis*; *Sa*, *Staphylococcus aureus*, *Ec*, *Escherichia coli*, *Af*, *Archaeoglobis fulgidus*). Those sequences for which crystal structures are available are highlighted with a Protein

Databank (PDB) accession number (in *red*). The subject of this work are the sequences associated with SSN cluster 2.

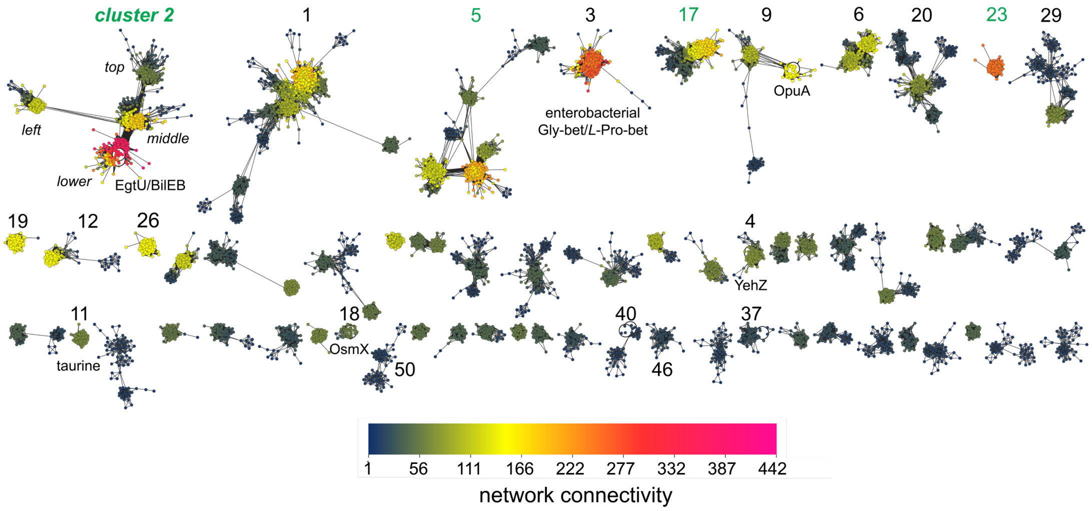


**Supplementary Figure 12. Network connectivity analysis**. Network connectivity representation for the top 58 SSN clusters (based on metanode cluster count). All clusters below this threshold are characterized by network connectivity values of ≤40 (see scale bar, *bottom*). The EgtU/BilEB subcluster in SSN cluster 2 (*lower*) along with SSN cluster 3 sequences encoding curated as enterobacterial (Gram-negative) glycine betaine/proline betaine SBPs, are among the most highly similar groups of sequences in the SSN database.

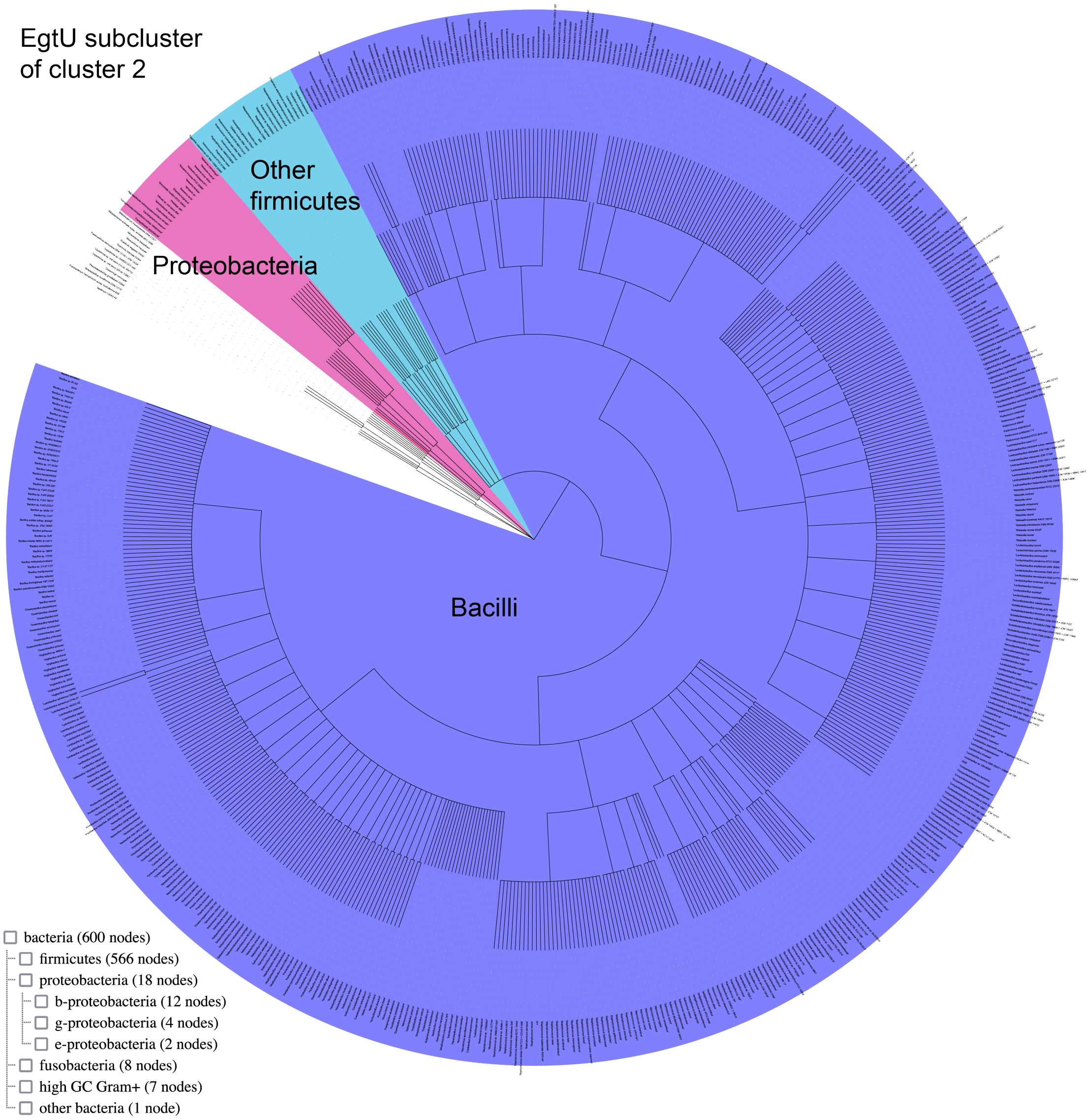


**Supplementary Figure 13. The biological range determined for the EgtU subcluster of cluster 2.** Taxonomy IDs extracted from the SSN analysis were supplied to the server NCBI Taxonomy common tree (https://www.ncbi.nlm.nih.gov/Taxonomy/CommonTree/wwwcmt.cgi)[2] to generate the phylogenetic tree, which was then visualized using iTOL.

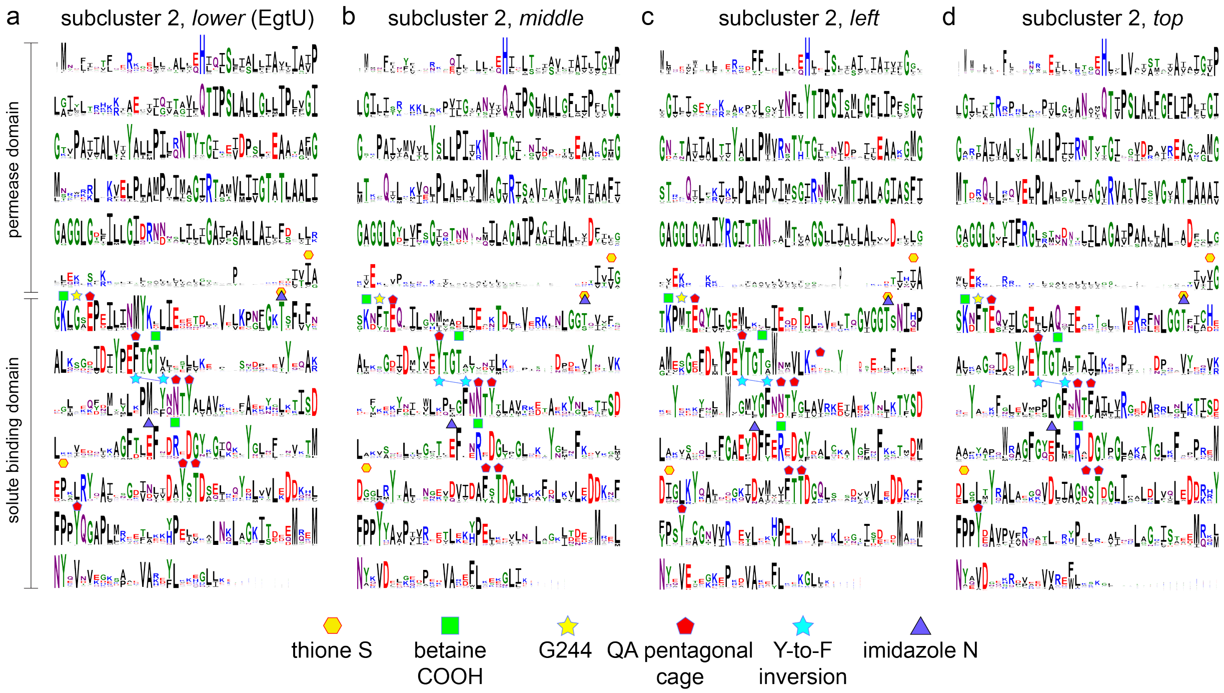


**Supplementary Figure 14. Sequence conservation maps of SSN cluster 2 subclusters.** Subclusters *lower* (*Sp*EgtU-like) (**a**), *middle* (*C. difficile* sequences)[3] (**b**), *left* (Streptococci, Clostridia, *H. pylori*) (**c**) and *upper* (Acidobacteria, Cyanobacteria) (**d**) multiple sequence alignments displayed as sequence logos[4]. Symbols (shown for reference) indicate conserved features of the ET binding site identified in the *Sp*EgtU subcluster of SSN cluster 2 (panel **a**) (see also Fig. 6c, main text), many of which are not broadly conserved in the other SSN cluster 2 subclusters (panels **b-d**). This analysis suggests that these other subclusters possess a QAC ligand specificity profile that is distinct from that of EgtU.

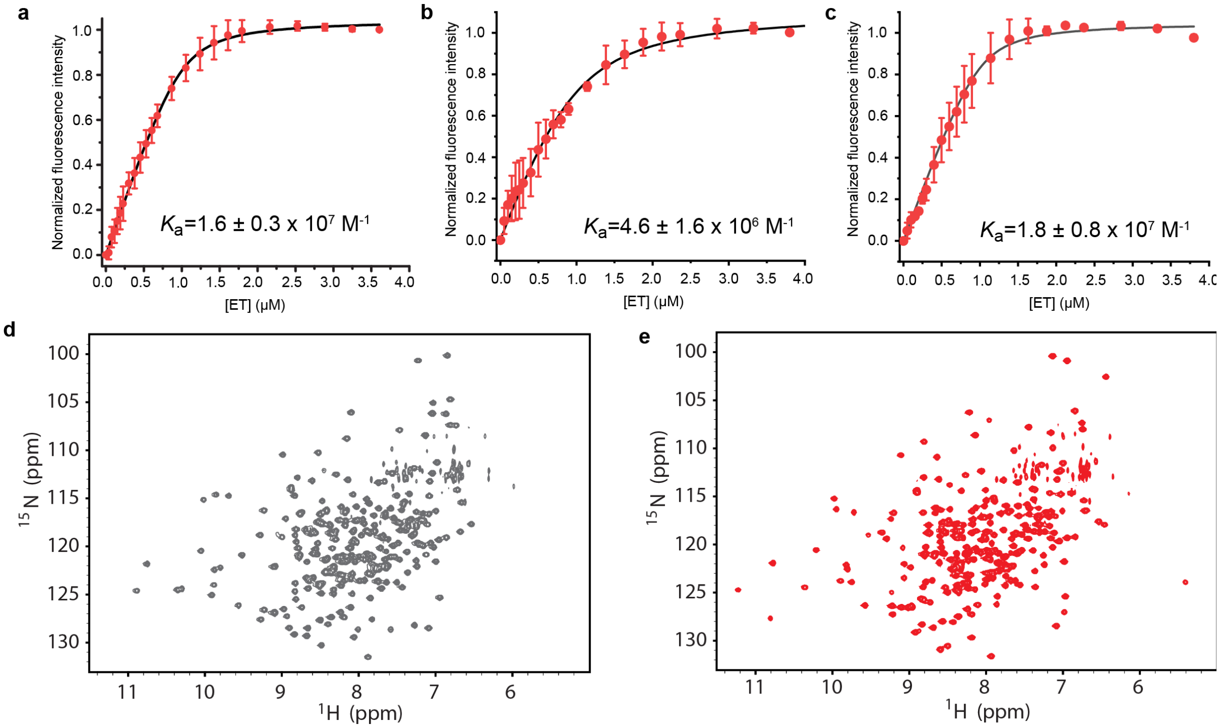


**Supplementary Figure 15. ET-binding properties of candidate EgtU homologs from other firmicutes. a-c,** Results of triplicate tyrosine fluorescence titrations of 1.0 μM *Ef*EgtU SBD (**a**), 1.0 μM *Sa*EgtU SBD (**b**) and 1.0 μM *Lm*EgtU SBD (**c**) with ET. The continuous line through the data shows the results of a fit to a 1:1 binding model with the $K_a$ indicated using DynaFit[5] with the parameters compiled in Table 1, main text. **d,** $^1$H,$^{15}$N TROSY spectrum of *Ef*EgtU SBD. **e,** $^1$H,$^{15}$N TROSY spectrum of *Ef*EgtU SBD in complex with equimolar ET.

### SUPPLEMENTARY TABLES

**Supplementary Table 1. Data collection and refinement statistics**

| | *Sp*EgtU SBD$_{CTT}$ | *Sp*EgtU SBD |
|---|---|---|
| *Data collection* | | |
| Wavelength (Å) | 1.07216 | 1.07216 |
| Space group | F222 | F222 |
| *Cell dimensions* | | |
| a, b, c (Å) | 118.00 129.02 206.91 | 119.87 128.74 207.54 |
| a, b, g (°) | 90.00 90.00 90.00 | 90.00 90.00 90.00 |
| Resolution * (Å) | 43.54 – 1.78 (1.82 – 1.78) | 43.86 – 2.44 (2.54 – 2.44) |
| $R_{sym}$ | 0.081 (0.983) | 0.167 (1.086) |
| $R_{meas}$ | 0.087 (1.062) | 0.195 (1.282) |
| $R_{pim}$ | 0.033 (0.400) | 0.073 (0.496) |
| Total reflections | 524963 (29990) | 211621 (22388) |
| No. unique reflections | 75117 (4251) | 29967 (3349) |
| CC1/2 | 0.998 (0.824) | 0.994 (0.714) |
| I/s(I) | 13.7 (1.6) | 8.5 (1.6) |
| Completeness (%) | 99.9 (100.0) | 99.9 (99.9) |
| Multiplicity | 7.0 (7.1) | 7.1 (6.7) |
| Wilson B-factor | 25.70 | 38.59 |
| *Refinement* | | |
| Resolution (Å) | 43.54 – 1.78 | 40.4 – 2.44 |
| No. unique reflections | 75056 (7462) | 29938 (2942) |
| $R_{work}$ | 0.1586 (0.2357) | 0.1834 (0.2696) |
| $R_{free}$ | 0.1863 (0.2671) | 0.2402 (0.3417) |
| *R.m.s.d values* | | |
| Bond lengths (Å) | 0.007 | 0.004 |
| Bond angles (°) | 0.96 | 0.603 |
| *No. atoms* | | |
| Protein | 4256 | 4324 |
| Ligand/ions | 115 | 70 |
| solvent | 539 | 156 |
| *B-factors* (Å$^2$) | | |
| Protein | 34.67 | 48.79 |

| | | |
|---|---|---|
| Ligand/ions | 47.85 | 42.07 |
| solvent | 40.92 | 41.49 |
| *Ramachandran plot* | | |
| Favored (%) | 98.0 | 98.0 |
| Allowed (%) | 1.6 | 1.6 |
| Outliers (%) | 0.4 | 0.4 |
| Clashscore | 3.25 | 7.21 |
| Rotamer outliers (%) | 0.2 | 1.3 |
| PDB code | 7TXL | 7TXK |

*Highest-resolution shell values are shown in parentheses.

**Supplementary Table 2. List of water molecules and associated B-factors in the structures of *Sp*EgtU SBD$_{CTT}$ and *Sp*EgtU SBD[a]**

| *Sp*EgtU SBD$_{CTT}$ | | | | *Sp*EgtU SBD | | | |
|---|---|---|---|---|---|---|---|
| Ori ID# | New ID# | Occupancy | B factor | Ori ID# | New ID# | Occupancy | B factor |
| HOH 23 | W1 | 1 | 24.73 | HOH 5 | W1 | 1 | 31.20 |
| HOH 22 | W2 | 1 | 24.32 | HOH 8 | W2 | 1 | 24.97 |
| HOH 33 | W3 | 1 | 23.90 | N/A | W3 | N/A | N/A |
| HOH 20 | W4 | 1 | 23.92 | HOH 49 | W4 | 1 | 31.52 |
| HOH 10 | W5 | 1 | 22.78 | HOH 32 | W5 | 1 | 27.79 |
| HOH 11 | W6 | 1 | 22.51 | HOH 60 | W6 | 1 | 29.67 |
| HOH 32 | W7 | 1 | 23.24 | HOH 43 | W7 | 1 | 34.67 |
| HOH 6 | W8 | 1 | 21.51 | HOH 4 | W8 | 1 | 38.00 |
| HOH 25 | W9 | 1 | 22.03 | HOH 51 | W9 | 1 | 34.67 |
| HOH 35 | | 1 | 26.39 | HOH 68 | | 1 | 41.48 |
| HOH 59 | | 1 | 24.77 | HOH 133 | | 1 | 38.62 |
| HOH 110 | | 1 | 30.93 | HOH 79 | | 1 | 41.05 |
| HOH 317 | | 1 | 42.01 | HOH 116 | | 1 | 49.28 |
| HOH 66 | | 1 | 25.63 | HOH 104 | | 1 | 40.73 |
| HOH 356 | | 1 | 35.41 | HOH 96 | | 1 | 35.69 |
| HOH 183 | | 1 | 36.34 | | | | |
| HOH 255 | | 1 | 40.86 | | | | |
| HOH 79 | | 1 | 29.09 | | | | |
| HOH 102 | | 1 | 29.60 | | | | |
| HOH 58 | | 1 | 26.19 | | | | |
| HOH 5 | | 1 | 22.78 | | | | |

[a]From PDB entries 7TXL and 7TXK, respectively. Water molecules 1-9 (W1-W9) correspond to those shown in Fig. 4, main text.

**Supplementary Table 3. List of *Streptococcus pneumoniae* D39 strains in this work**

| Strain ID | Genotype | Ref |
|---|---|---|
| IU1781 | D39 *rpsL1* (WT) | [6] |
| IU18405 | D39 *rpsL1* Δ*spd_1642,* markerless (Δ*egtU*$^{ML}$) | This work |
| IU18789 | D39 *rpsL1* Δ*spd_1642,* markerless, repaired (Δ*egtU*$^{ML}$ repaired) | This work |

**Supplementary Table 4. Primers used in this study[a]**

| Primer IDs | Sequence 5' to 3' |
|---|---|
| ***Sp*EgtU mutant strains** | |
| 1642_outside_FP | GGAACTGCGACTTCTGCGC |
| 1642_outside_RP | CTCCCAGACTGTTTCACTCCCG |
| 1642_KanrpsL_NT_RP | CATTATCCATTAAAAATCAAACGGATCCTAATCCTGAAAAGTTGCAATTAAATTAGTC |
| 1642_KanrpsL_CT_FP | CAAAAGCATAAGGAAAGGGGCCCCTCCAAGAACAAGGTTTGTTGAAGAAATGATGG |
| 1642_ML_Rev | CCTTGTTCTTGGAGATCCTGAAAAGTTGCAATTAAATTAGTCATGAATACTACC |
| 1642_ML_Fwd | GCAACTTTTCAGGATCTCCAAGAACAAGGTTTGTTGAAGAAATGATG |
| **Protein expression** | |
| *Sp*EgtU-SBD_pSUMO_FP | GAACAGATTGGATCCCATATGGAGAAGGAAAACTTGATTATTGCTGGGAAAATAGGC |
| *Sp*EgtU-SBD_pSUMO_RP | GTGCTCGAGTGCGGCCGCAAGCTTTCATTTCTTCAACAAACCTTGTTCTTGGAGAAACTC |
| *Sp*EgtU-SBD_Y419F_FP | GGGGATATTCAAATCACGGATGCCTTTTCGACTGATGCGGAATTGGAGC |
| *Sp*EgtU-SBD_Y419F_RP | GCTCCAATTCCGCATCAGTCGAAAAGGCATCCGTGATTTGAATATCCCC |
| *Sp*EgtU-SBD_G244F_FP | GGAAAACTTGATTATTGCTGGGAAAATATTCCCAGAACCAGAAATTTTGGCTAATATG |
| *Sp*EgtU-SBD_G244F_RP | CATATTAGCCAAAATTTCTGGTTCTGGGAATATTTTCCCAGCAATAATCAAGTTTTCC |
| *Sp*EgtU-SBD_F277W_FP | GTTAAACCGAATTTTGGGACGACAAGTTGGCTTTATGAAGCTCTGAAAAAAGGTG |
| *Sp*EgtU-SBD_F277W_RP | CACCTTTTTTCAGAGCTTCATAAAGCCAACTTGTCGTCCCAAAATTCGGTTTAAC |
| *Sp*EgtU-SBD_L374C_FP | GCAGCTGAAGGCAGGCTTTACGTGTGAGTTTAACGACCGTGAAGATGGAAATAAGG |
| *Sp*EgtU-SBD_L374C_RP | CCTTATTTCCATCTTCACGGTCGTTAAACTCACACGTAAAGCCTGCCTTCAGCTGC |
| *Sp*EgtU-SBD_F293Y_FP | GAAAAAAGGTGATATTGACATTTATCCTGAATATACCGGTACGGTGACTGAAAG |
| *Sp*EgtU-SBD_F293Y_RP | CTTTCAGTCACCGTACCGGTATATTCAGGATAAATGTCAATATCACCTTTTTTC |
| SpEgtU-SBD_CTT_RP | GTGCGGCCGCAAGCTTTCATTTCTTCAACAAACCTGTTCTTGGAGAAACTCCTTGGCTA |
| *Ef*EgtU-SBD_pSUMO_FP | GAACAGATTGGAGGATCCCATATGAAGGAAAAACAGCTGACAATTGCTGGC |

| | |
|---|---|
| *Ef*EgtU-SBD_pSUMO_RP | TCGAGTGCGGCCGCAAGCTTttaTTTAAGAAGATTCTTTTCTTTTAAATAGTCTTTCGCC |
| *Sp*EgtU-S_P1 | TGGTTCTGGTGATGGTTGAAGCAAACTTTCAGTCACCG |
| *Sp*EgtU-S_P2 | GAATACAACTTTCCTCCTCCCAAGGTGAGTCATGAGCCAG |
| *Sp*EgtU-S_P3 | GCTTCAACCATCACCAGAACCAAACGTATACATCACTGCTGACAAACAAAAAAAACGG |
| *Sp*EgtU-S_P4 | CATACTACCACCTGTACCACCCTTATAAAGCTCATCCATGCCGTGAGTGATACC |
| *Sp*EgtU-S_P5 | GGGTGGTACAGGTGGTAGTATGAAAGGAGAAGAGCTGTTCACAGGTG |
| *Sp*EgtU-S_P6 | CACCTTGGGAGGAGGAAAGTTGTATTCAAGTTTGTGTCCAAGGATGTTTCC |
| *Sa*EgtU-SBD_pSUMO_FP | GAACAGATTGGAGGATCCCATATGGGTGATAAAATTACGTTAGCTGGAAAGCTTGG |
| *Sa*EgtU-SBD_pSUMO_RP | GCTCGAGTGCGGCCGCAAGCTTTTATTTGATTAACCCTTTTGCTTTTAAATAATCTTTCG |
| *Lm*EgtU-SBD_pSUMO_FP | AGATTGGAGGATCCCATATGTCGGATAAAAAGGAAATTACAATTGCTGGTAAATTAGG |
| *Lm*EgtU-SBD_pSUMO_RP | CTCGAGTGCGGCCGCAAGCTTTATTTAATAATACCTTGATCTTTCAAATAGTCTTTGGC |

[a]All sequences are written 5' to 3'. FP, forward primer; RP, reverse primer.

**Supplementary Table 5. Molar extinction coefficients of purified proteins at 280 nm ($\varepsilon_{280}$) used in this work**

| Protein | Mutation | MW, Da | molar extinction coefficient ($\varepsilon_{280}$)[a], $M^{-1}$ $cm^{-1}$ |
|---|---|---|---|
| *Sp*EgtU SBD | WT | 31032.4 | 20,860 |
| | CTT[b] | 30695.0 | 20,860 |
| | Y419F | 31016.4 | 19,370 |
| | G244F | 31122.5 | 20,860 |
| | F293Y | 31048.4 | 22,350 |
| | F227W/L374C | 31061.4 | 26,360 |
| SpEgtU SBD-sfGFP | – | 58508.4 | 41,260 |
| *Ef*EgtU SBD | WT | 31373.7 | 22,350 |
| *Sa*EgtU SBD | WT | 31638.6 | 22,350 |
| *Lm*EgtU SBD[c] | WT | 31351.8 | 23,840 |

[a]Assuming that all Cys residues are in the reduced form. [b]CTT, C-terminally truncated. [c]Also known as BilEB[7].

### SUPPLEMENTARY REFERENCES


1. Zallot, R., Oberg, N. & Gerlt, J.A. The EFI Web Resource for Genomic Enzymology Tools: Leveraging Protein, Genome, and Metagenome Databases to Discover Novel Enzymes and Metabolic Pathways. *Biochemistry* **58**, 4169-4182 (2019).
2. Letunic, I. & Bork, P. Interactive Tree Of Life (iTOL) v5: An online tool for phylogenetic tree display and annotation. *Nucleic Acids Res* **49**, W293-W296 (2021).

3. Michel, A.M. et al. Cellular adaptation of *Clostridioides difficile* to high salinity encompasses a compatible solute-responsive change in cell morphology. *Environ Microbiol* **24**, 1499-1517 (2022).
4. Crooks, G.E., Hon, G., Chandonia, J.M. & Brenner, S.E. WebLogo: a sequence logo generator. *Genome Res* **14**, 1188-90 (2004).
5. Kuzmic, P. Program DYNAFIT for the analysis of enzyme kinetic data: application to HIV proteinase. *Anal Biochem* **237**, 260-273 (1996).
6. Lanie, J.A. et al. Genome sequence of Avery's virulent serotype 2 strain D39 of *Streptococcus pneumoniae* and comparison with that of unencapsulated laboratory strain R6. *J Bacteriol* **189**, 38-51 (2007).
7. Ruiz, S.J., Schuurman-Wolters, G.K. & Poolman, B. Crystal structure of the substrate-binding domain from *Listeria monocytogenes* bile-resistance determinant BilE. *Crystals* **6**, 162 (2016).
